# Aneuploidy in human embryos is associated with a maternal age-independent increase in mitochondrial DNA content and an enrichment of ultra-rare mitochondrial DNA variants

**DOI:** 10.1101/2022.10.14.512116

**Authors:** Maxim Ri, Natalia Ree, Sergey Oreshkov, Maria Tofilo, Irina Zvereva, Anastasia Kirillova, Konstantin Gunbin, Valerian Yurov, Andres Salumets, Dori C. Woods, Konstantin Khrapko, Jacques Fellay, Jonathan L. Tilly, Ilya Mazunin, Konstantin Popadin

## Abstract

Using low-coverage, whole-genome sequences of trophectoderm biopsies from 11,610 human blastocyst-stage embryos, we analyzed the relationship between chromosomal abnormalities and mitochondrial (mt) DNA dynamics. Comparing 6,208 aneuploid and 5,402 euploid embryos in cohort studies, we found that mtDNA content in aneuploid embryos was significantly higher than that in euploid embryos. This outcome was confirmed through intrafamilial analyses of embryos with matched parents and *in vitro* fertilization cycles, and it occurred independent of maternal age. Additional human population-based studies uncovered a higher abundance of ultra-rare mtDNA variants located in never-altered positions in the human population in aneuploid compared to euploid embryos in both cohort- and family-based analyses. This maternal age-independent association of increased mtDNA content and aneuploidy in human embryos may reflect a novel mechanism of purifying selection against potentially deleterious mtDNA variants, which arise from germline or early developmental mtDNA damaging events, that occurs in human embryos prior to implantation.

## MAIN

Coordination of bioenergetics by mitochondria is central to the function of most cells, and thus a broad spectrum of processes that require ATP are negatively impacted if mitochondrial function becomes impaired. Mitochondria also have the unique feature of being controlled by two distinct genomes, with one localized in the nucleus and the other localized within mitochondria. The mitochondrial genome, referred to as mitochondrial DNA (mtDNA), is comprised of 37 genes that code for 22 transfer RNAs and 2 ribosomal RNAs, as well as for 13 proteins which are building blocks for the electron transport chain. An important aspect of mtDNA is that its mutational rate is estimated to be at least one log-order higher than that of the nuclear genome^1^, which reflects the proximity of mtDNA to reactive oxygen species (ROS) generated along the inner mitochondrial membrane during ATP synthesis as well as a limited capacity of cells to repair mutations. Impairments in mitochondrial function can therefore arise intrinsically from defects in mitochondrial gene expression as well as extrinsically from defects in expression of nuclear genes that influence mitochondrial dynamics. Accordingly, there is tremendous interest in defining the contribution of mitochondrial dysfunction and mtDNA mutations in the etiology of diseases, such as cancer, MELAS (mitochondrial encephalopathy lactic acidosis with stroke-like episodes) and MERRF (myoclonic epilepsy ragged red fibers), as well as in aging-associated organ dysfunction and failure^2,3^.

The fundamental importance of mitochondria to organismal development, survival and inheritance is particularly well-illustrated by studies of human development, which have established a strong causative association of impaired mitochondrial function with chromosomal abnormalities in eggs and embryos that lead to infertility, embryo implantation failure, and neonatal disorders^4–6^. For example, studies of mouse and human oocytes have shown that meiotic spindle abnormalities, embryonic developmental arrest, and failed conception are tied to perturbed mitochondrial function and reduced ATP levels^7–9^. Clinical studies have demonstrated that mtDNA content is elevated in trophectoderm biopsies of blastocysts from women aged 38 years and older compared to blastocysts from women 37 years of age or younger^10^. When chromosomally abnormal blastocysts were analyzed, a tight association between mtDNA content and aneuploidy was detected independent of maternal age^10^. Subsequent blinded clinical studies reaffirmed an association between elevated mtDNA content in trophectoderm biopsies and adverse pregnancy outcomes.^11,12^ However, the robustness of this association has been heavily debated based on conflicting data from others^13,14^, which may reflect methodological differences and low sample sizes across the various studies on this topic^15^.

In an effort to resolve these questions and further explore mtDNA dynamics during early embryonic development, data from 11,610 human blastocyst-stage embryos were used to evaluate associations between chromosomal abnormalities and mtDNA (Fig. 1). For each embryo, 5–10 trophectoderm cells with no signs of degradation were utilized for preimplantation genetic testing for aneuploidies (PGT-A) via two different analytical platforms: Illumina VeriSeq PGS (*n* = 3,657 embryos) and AB Vector AB-PGT (*n* = 7,953 embryos). Low-coverage, whole-genome sequence (WGS) data obtained from trophectoderm biopsies were used to guide clinical decisions on whether a given embryo was recommended for transfer (RFT; *n* = 5,402 euploid embryos or embryos with a low level of mosaicism: 1,648 from VeriSeq PGS and 3,754 from AB-PGT) or not recommended for transfer (NRFT; *n* = 6,208 aneuploid embryos or embryos with a high level of mosaicism: 2,009 from VeriSeq PGS and 4,199 from AB-PGT). For each embryo, we derived values for mtDNA content by using the number of reads mapped to the mitochondrial genome normalized to the number of reads mapped to the nuclear genome.

**Fig. 1.**
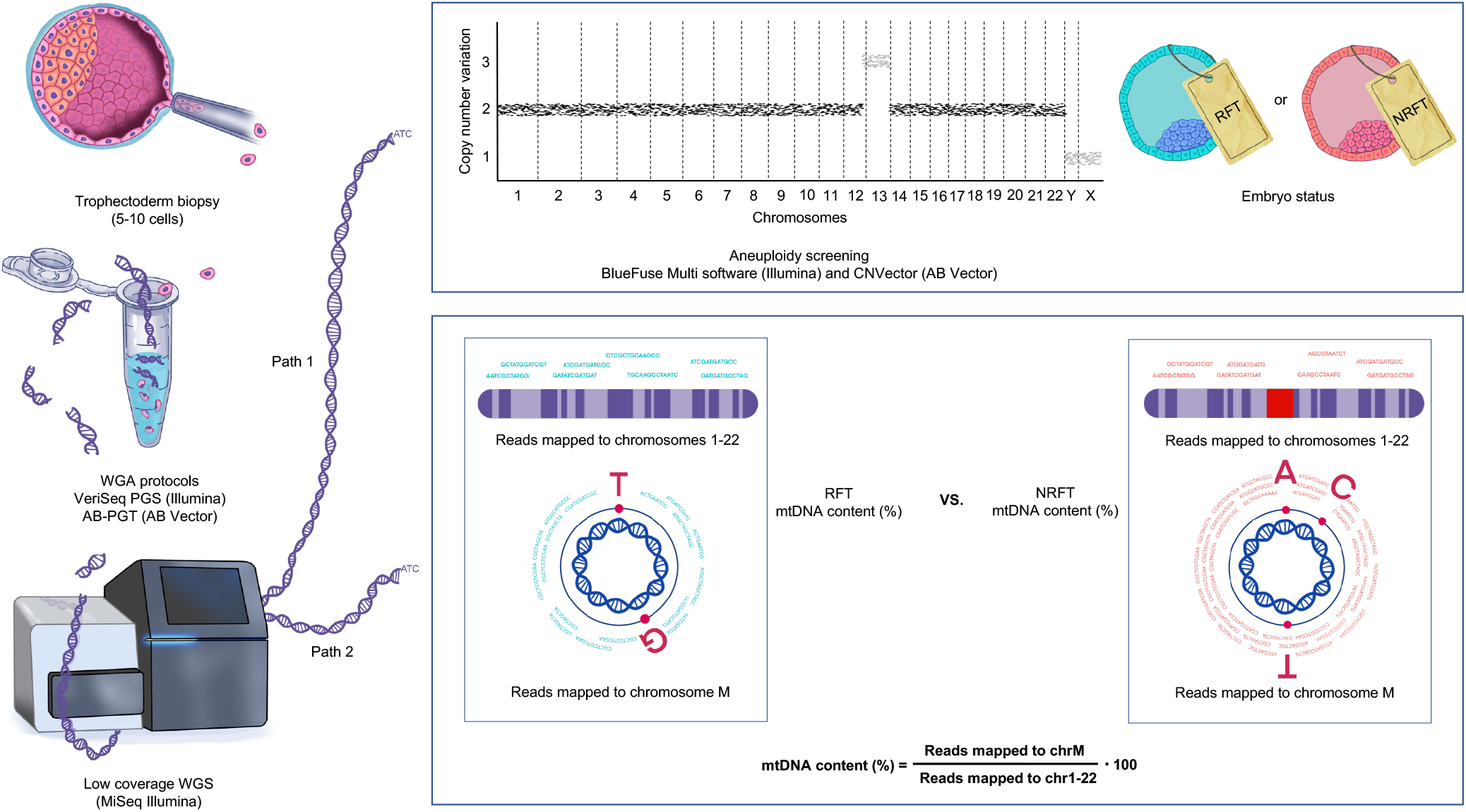
Overview of the study design and workflow. We performed low coverage, whole genome sequence (WGS) analysis of trophectoderm biopsies from 11,610 morphologically good quality human embryos as a preimplantation diagnostic for aneuploidy. Path 1: the chromosomal status of each embryo was annotated as either euploid (recommended for transfer, RFT) or aneuploid (not recommended for transfer, NRFT) using BlueFuse Multi software (Illumina) and CNVector (AB Vector) tools. The clinical report was based on the analysis of CNV charts (see Methods). Path 2: the mitochondrial component of the WGS was analyzed in order to assess mitochondrial DNA (mtDNA) content and the presence of ultra-rare-potentially deleterious mtDNA variants which are marked by red dots.

Using this analytical workflow (Fig. 1), we observed that mtDNA content was significantly higher in NRFT versus RFT embryos, irrespective of which platform was employed for the analysis (VeriSeq PGS: nominal *P*-value = 1.70E-15; corrected *P*-value = 0.0008; AB-PGT: nominal *P*-value = 2.25E-12; corrected *P*-value = 0.0262) (Fig. 2a, Supplementary Data 1). By splitting the entire mitochondrial genome into three separate regions—major arc, minor arc, and origins of replication, we demonstrated the uniformity of this observation in that mtDNA content was elevated in NRFT versus RFT embryos for all three regions (Supplementary Data 2). A multiple logistic regression model was then performed to test interrelationships between maternal age, mtDNA content, and aneuploidy. We observed that advanced maternal age and increased mtDNA content were significantly and independently associated with a high risk of aneuploidies (Supplementary Data 3–7). Additional analyses confirmed an increase in mtDNA content in NRFT embryos irrespective of total coverage of the genome (Supplementary Data 8), sex chromosome karyotype, type of chromosomal abnormality, embryo developmental day, or blastocyst morphology (Supplementary Data 9-10). To control for parental age and other family-related factors, such as nuclear genetics of both parents, maternal mtDNA haplogroups, social-economical status, and dietary differences, we next analyzed a total of 1,827 families in our dataset that had at least one RFT embryo and one NRFT embryo obtained from the same controlled ovarian stimulation and *in vitro* fertilization (IVF) cycle (Fig. 2b). This intrafamilial pairwise analysis confirmed a significant increase in the mean mtDNA content in NRFT versus RFT embryos of common parental origin and IVF cycle (6.7% increase with AB-PGT, 18.1% increase with VeriSeq PGS; Fig. 2b; Supplementary Data 11), thus verifying the main result obtained at the cohort level (Fig. 2a).

**Fig. 2.**
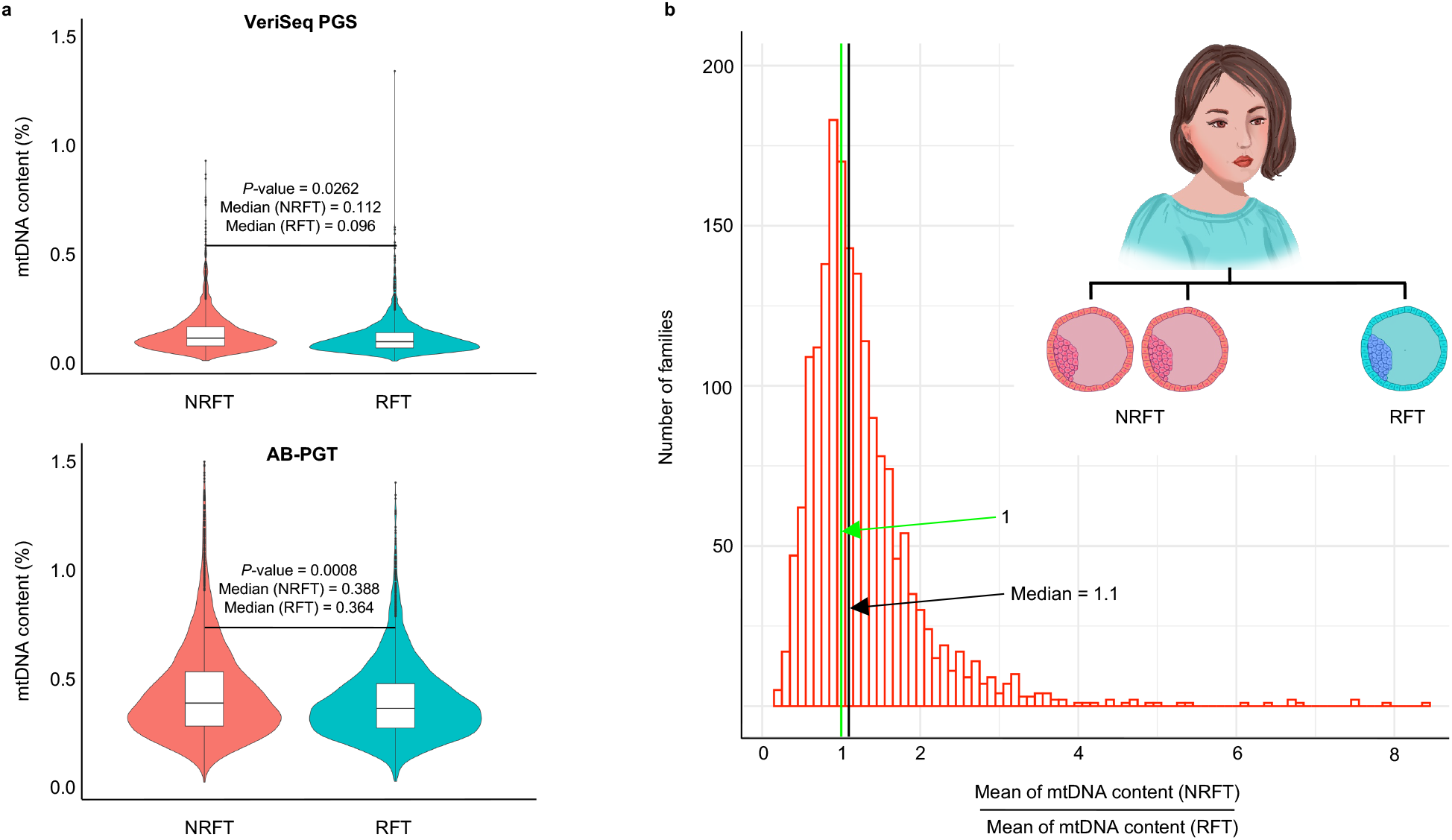
NRFT embryos have a higher mtDNA content compared to RFT embryos. **a,** Cohort-based analysis of 11,610 embryos shows that mtDNA content was significantly higher in NRFT versus RFT embryos irrespective of the platform utilized: for VeriSeq PGS, medians were 0.112 and 0.096 for NRFT and RFT embryos, respectively (nominal *P*-value=1.70E-15, Mann-Whitney U test; corrected *P*-value = 0.0008); for AB-PGT, medians were 0.388 and 0.364 for NRFT and RFT embryos, respectively (nominal *P*-value=2.25E-12, Mann-Whitney U test; corrected *P*-value = 0.0262). Additional analyses confirmed this observation and its independence from maternal age (see Supplementary Data 2–8). **b,** Familial-based analysis of embryos using a pairwise comparison of mtDNA content between NRFT and RFT embryos within the same family (*n* = 1,827 families, with VeriSeq PGS and AB-PGT platforms analyzed together) confirmed a significant increase in the mean mtDNA content in NRFT versus RFT embryos of the same family (*P*-value = 3.56E-36, paired Mann-Whitney U test; see also Supplementary Data 11).

The observed increase in mtDNA content in aneuploid embryos (Fig. 2a,b) could be either a consequence or a cause of aneuploidies. One plausible causative scenario entails a deleterious mitochondrial or nuclear-encoded DNA variant of either germline or early somatic origin that perturbs mitochondrial function, which then triggers a compensatory mechanism involving mtDNA overreplication^16^. However, the increase in mtDNA does not resolve the mitochondrial dysfunction, which compromises ATP generation leading to chromosomal abnormalities and implantation failure^7–9^. To test this possibility, we focused on mtDNA variants, which could be called in some regions of moderately high coverage. Similar regions were absent in the nuclear genome. Using all alternative mtDNA variants supported by at least one forward and one backward read (Fig. 3a), we tested whether there was an excess of potentially deleterious variants in cohorts of NRFT versus RFT embryos (Supplementary Data 12). As a proxy readout for the deleterious potential of each altered mtDNA position, we used corresponding frequencies of human population mtDNA polymorphisms based on 195,983 completely-sequenced mitochondrial genomes from around the globe^17^. We observed that frequently-polymorphic (with three or more homoplasmic carriers of an alternative allele in a given position) and rarely-polymorphic (with at least one carrier of a heteroplasmic alternative variant and less than three carriers of homoplasmic variants) positions were uniformly distributed among RFT and NRFT embryos. In contrast, never-altered positions (positions without alternative variants) were enriched in NRFT embryos (Supplementary Data 12). Additional analyses, taking into consideration the nucleotide coverage and recurrence of mtDNA mutations among NRFT and RFT embryos, confirmed an excess of these ultra-rare mtDNA mutations in NRFT embryos (Supplementary Data 12).

**Fig. 3.**
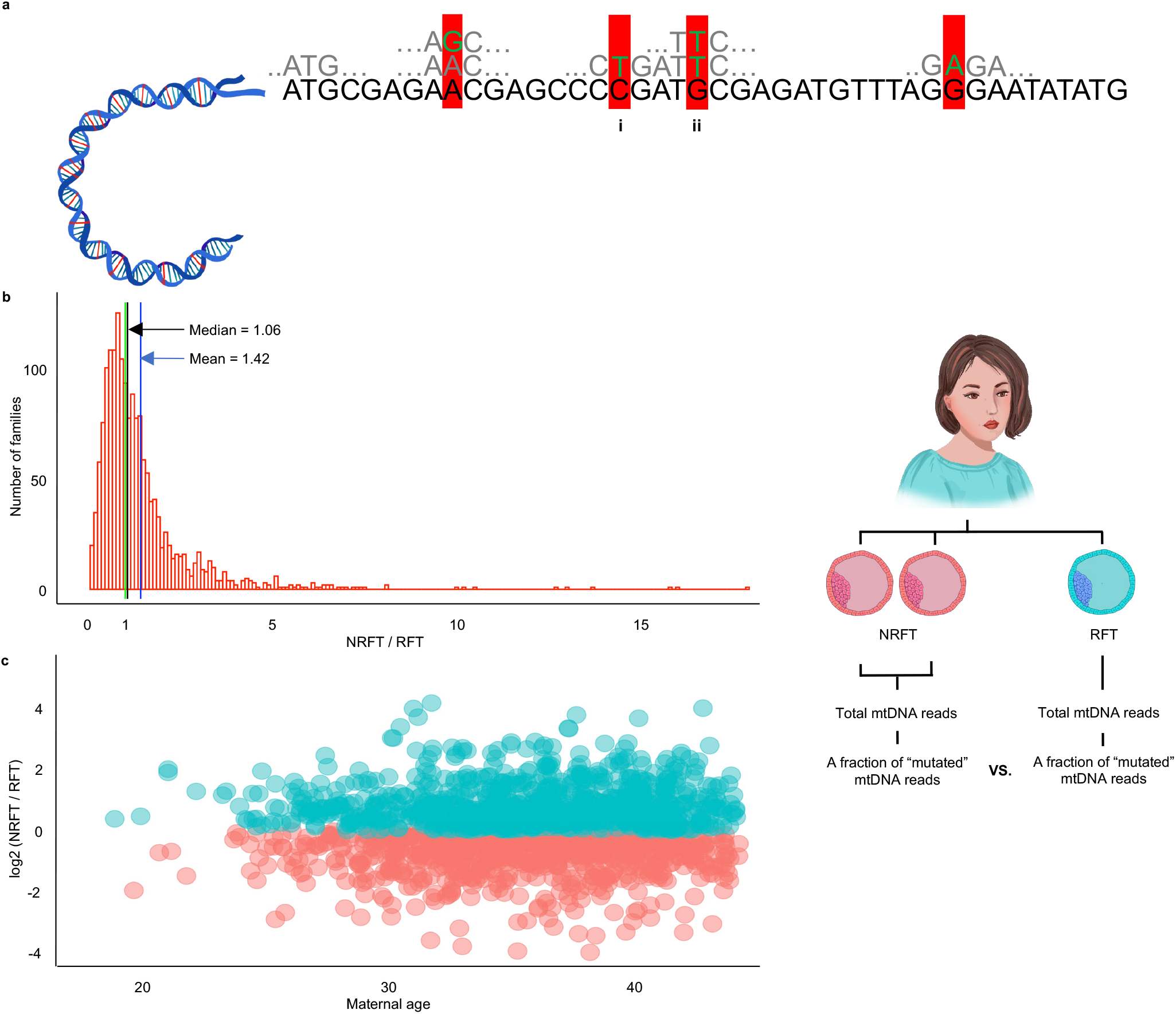
Ultra-rare mtDNA variants are enriched in NRFT versus RFT embryos: familybased approach. **a,** A scheme of the analysis. Positions in the mitochondrial genome that are never altered in the HELIX database (n = 6,316) are marked by red background. Nucleotides that deviate from the reference alleles in the never-altered positions are marked in green. Only variants supported by at least one forward and one backward read (ii in the scheme) were used for the analysis. **b**, Each family used in the analysis had at least one NRFT embryo and at least one RFT embryo. A fraction of “mutated” mtDNA reads, i.e. reads having an alternative allele on never-altered position, has been estimated separately for all NRFT embryos and RFT embryos. The ratio of fractions of the mutated reads in NRFT to RFT is plotted on the histogram (n = 1,686). Median and mean of the ratios significantly higher than the expected value of 1 (*P*-value < 2.2E-16, Wilcox test with mu equals 1). **c**, Log ratio of fractions of mutated reads in NFRT versus RFT for each family as a function of maternal age (n = 1,646). An excess of families with a ratio greater than 0 is uniformly distributed across different maternal ages. See Supplementary Data 13 for details.

Next, we performed our analysis of ultra-rare mtDNA variants on the familial level by focusing on 1,827 families that had at least one NRFT embryo and at least one RFT embryo. In order to maximize coverage for this analysis, all mtDNA reads from NRFT embryos and, separately, all mtDNA reads from RFT embryos within each family were merged to create a single dataset each for NRFT and RFT mtDNA reads per family analyzed. Next, the fraction of mutated mtDNA reads having an alternative allele on a never-altered position was assessed using the NRFT and RFT datasets within the same family. The expectation that the fraction of mutated reads is the same among NRFT and RFT datasets within each family was rejected, since the ratios of the mutated reads in NRFT to RFT datasets had a median significantly higher than one (Fig. 3, Supplementary Data 13; *P*-value < 2E-16 using the Wilcoxon test with the theoretical mean value = 1). Overall, our family-based analysis also showed an excess of ultra-rare variants in NRFT versus RFT embryos, thus confirming the results obtained on the cohort level (Supplementary Data 12).

Past studies have proposed that embryonic aneuploidies could also arise in tandem with or as a consequence of the accumulation of mtDNA defects within oocytes as females age^5,18^, similar to what has been observed in somatic postmitotic tissues^19^ (Fig. 4). However, this model predicts a strong association of mitochondrial dysfunction and mtDNA overreplication with advanced maternal age, which was not observed in our study. Thus, an early mitochondrial damage scenario with rapid and strong effects, as opposed to a slow and progressive accumulation of mtDNA mutations in oocytes, appears to be a more reasonable explanation for maternal age-independent aneuploidies detected in human embryos (Fig. 4). There are several ways that mitochondrial phenotype and function could be perturbed fast in the absence of predisposing genetic variants from the previous generation: (a) de-novo germline mutations in nuclear-encoded mitochondrial regulatory genes; (b) de-novo or inherited low-heteroplasmic germline mtDNA variant, which after narrow stochastic mtDNA bottleneck of about 20 inherited mtDNA copies can increase its heteroplasmy level drastically by chance; (c) deterministic positive selection of deleterious mtDNA variants in the germline^20^; (d) mtDNA mutations with dominant negative effects on mitochondrial phenotype and function even at very low heteroplasmy levels^21^; and, (e) damage of oocytes in early life by chemical or hormonal stresses that lead to mtDNA nucleotide modifications, such as N-deoxyadenosine methylation, which have pleiotropic effects on the mitochondrion^22^. As described above, the ultra-rare mtDNA variants enriched in NRFT embryos (Fig. 3, Fig. 4) can assigned to categories b, c, d. Irrespective of the initiating event, it has been proposed that deleterious mtDNA variants are selected out by the waves of apoptosis which occur naturally in oocytes during embryonic development^23,24^. Our data support an extension of this timeline for purifying selection in that some deleterious mitochondrial variants may also be selected out through aneuploidies that arise in embryos derived from the fertilization of those oocytes that survive apoptosis (Fig. 4).

**Fig. 4.**
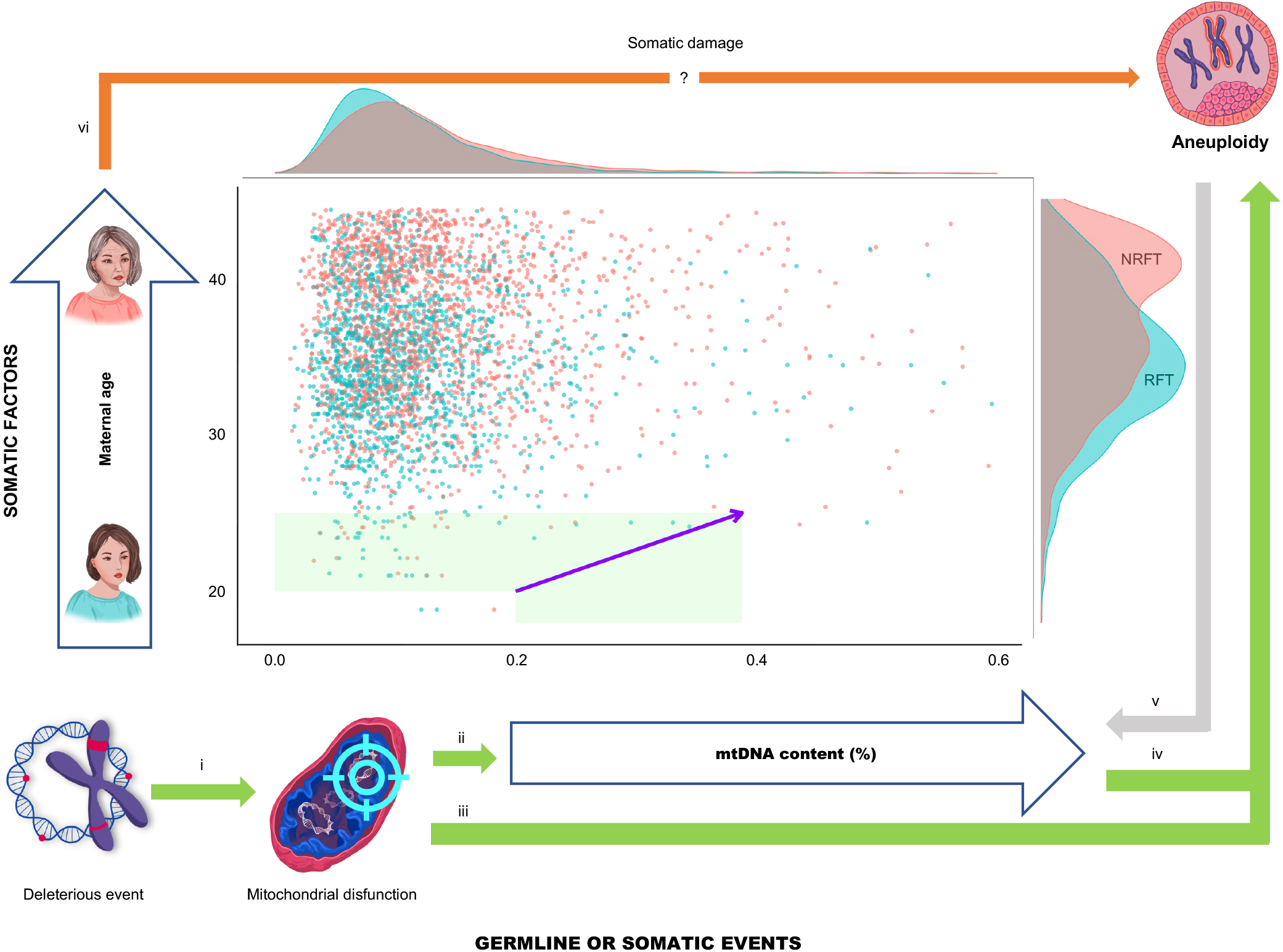
Schematic summary of potential causal relationships between early mitochondrial damage, mtDNA content and aneuploidy, and maternal age. The *x*-axis (mtDNA content) and *y*-axis (maternal age) are two independent factors that affect the risk of aneuploidies in human embryos (see Supplementary Data 3–8). NRFT embryos are marked by red points, and RFT embryos are marked by blue points. The purple arrow indicates an increase in aneuploidy risk: each additional 5 years of maternal age is equivalent to an increase in mtDNA content by 0.188% (see Supplementary Data 7). We interpret the increased mtDNA content, due to its age-independency (see Supplementary Data 4), as a germline or early somatic risk factor of aneuploidies. The proposed chain of events linking early mitochondrial damage to aneuploidy is as follows: (*arrow i*) a germline deleterious event, such as a genetic or epigenetic change, affects the nuclear or mitochondrial genome, leading to mitochondrial dysfunction; (*arrow ii*) in an effort to compensate for this dysfunction, cells overreplicate mtDNA; (*arrow iii*) ongoing mitochondrial dysfunction, even with mtDNA overreplication, is accompanied by ATP deficiency, which results in aneuploidy; (*arrow iv*) increased mtDNA content is a marker of aneuploidy, connecting the causative deleterious germline event with the aneuploidy. The relationship between mtDNA content and aneuploidy may be a consequence of aneuploidy itself (*arrow v*), where aneuploidy leads to an increase in mtDNA content as a cellular response. The *y*-axis represents a combination of all somatic factors associated with maternal aging, such as the accumulation of somatic mutations, epigenetic changes, and misfolded proteins, all of which can cause aneuploidy (*arrow vi*). The scatterplot used in this summary is from Supplementary Data 5.

In summary, the large sample size studied, combined with the family-based pairwise embryo analysis, has allowed us to unambiguously confirm an association of elevated mtDNA content with aneuploidy in human embryos, as well as to generate the first evidence for ultra-rare, and potentially-deleterious, mtDNA variants in driving mtDNA overreplication associated with aneuploidy. We should emphasize that the clinical outcomes of the embryo transfers were not assessed in this study. However, analysis of mtDNA content among euploid human embryos has demonstrated that increased mtDNA content is associated with a low viability score^25^, suggesting that the clinical applications of assessing mtDNA content can be rather broad. Here we have demonstrated that mtDNA content can be used to predict aneuploidies (Supplementary Data 6). Thus, mtDNA content may serve as an additional marker of aneuploidies, which is especially important in borderline cases of mosaicism or noisy low-coverage nuclear DNA data. Considering the maternal age-independent association of mtDNA content with aneuploidies in human blastocysts, this parameter may be of special interest to younger mothers with familial cases of aneuploidies or mitochondrial diseases. To highlight the potential importance of increased mtDNA content in the prediction of aneuploidy risk for young mothers, we performed a decision tree analysis which demonstrated that mtDNA content is indeed the most important factor among young mothers and thus can be used to classify the risk of aneuploidies (Supplementary Data 14). The use of mtDNA content as an independent predictive marker of aneuploidy risk might also improve the sensitivity and precision of non-invasive preimplantation genetic testing for aneuploidies (niPGT-A), which is based on DNA extracted from the medium in which embryos are cultured^26^. It has been shown that this medium also contains mtDNA, and its level is biologically meaningful^27,28^. Finally, an additional application of trophectoderm biopsy mtDNA content analysis could be an improved annotation of triploid embryos of the female sex. Triploidy accounts for ~2% of natural pregnancies and 15% of cytogenetically abnormal miscarriages^29^. Commercial software, such as Bluefuse Multi Software (Illumina), allows for visualization of only triploid embryos with Y chromosomes, while the ability to call triploidy for female sex embryos requires additional costly procedures^30^. Using mtDNA content from standard PGT-A as a marker of aneuploidies, it may be possible to provide low-cost options for prediction of triploid female sex embryos.

## ETHICS STATEMENT

This study was approved by the Ethics Committee of Fomin Clinic, Moscow, Russia (ID No: 02/2021). Informed consent was obtained from all participants included in the study.

## ACKNOWLEDGEMENTS

Analysis of mitochondrial content by M.T., A.K. and I.M. was supported by the Russian Science Foundation (No. 21-75-10081). The analysis of the effect of maternal age on the risk of aneuploidies by K.G. was supported by Russian Science Foundation (No. 21-75-20145). Data analysis by K.K., D.C.W. and J.L.T was supported by the United States National Institutes of Health (NIH R01-HD091439). M.R. was supported by the Ministry of Science and Higher Education of the Russian Federation (No. 075-02-2022-872). The authors thank Kateryna Makova and Evgenii Tretiakov for valuable suggestions.

## AUTHOR CONTRIBUTIONS

I.M. and K.P. conceived and designed the study. A.K. and I.Z. conducted the culture and morphological assessment of embryos. M.T. and I.M. conducted PGT-A testing and divided the embryos into groups. M.R. developed a custom bioinformatics pipeline. M.R., K.P., S.O., K.G. and V.Y. conducted statistical analysis. M.R., N.R., J.L.T., I.M., and K.P. wrote the paper. N.R. illustrated figures in the main text. All authors discussed the results, offered interpretations of the data, provided input, and approved the final paper for submission.

## METHODS

### Study design

Patients using fresh or vitrified autologous oocytes undergoing assisted reproduction with PGT-A were included in the study, which was performed with IRB approval (ID 02/2021) at Fomin’s Clinic (Moscow, Russia). Embryos were cultured for 5–7 days to the blastocyst stage; all embryos were classified based on morphological criteria using the Gardner system^31^. For PGT-A analysis, 5–10 trophectoderm cells were biopsied from each viable blastocyst, and embryos were then either recommended for transfer or not recommended for transfer using the following workflow: FASTQ files were obtained from MiSeq Illumina based on two diagnostic tests (Illumina VeriSeq PGS and AB Vector AB-PGT) and aligned on GRCh37 by BlueFuse Multi software (Illumina) and CNVector tools (AB Vector). Clinical reports were based on CNVector chart analysis. Euploid embryos and embryos with mosaicism lower than 20% were recommended for transfer. Embryos with aneuploidy, deletions, duplications, mosaic aneuploidy of more than 80%, and mosaicism of three or more chromosomes were not recommended for transfer. Embryos with mosaic aneuploidies 13, 14, 16, 18, 21, and 45X were not recommended to transfer, regardless of the mosaicism percentage^32,33^. For embryos with a single mosaic aneuploidy between 20% - 80% of a chromosome not listed above, genetic counseling was recommended to assess the possible risk of transfer, and such embryos were filtered from our genome analysis (see below).

### Genome analysis

Raw reads from WGS were aligned to the human GRCh38 genome with circular mDNA using BWA (version 0.7.17), with aln and samse algorithms. The output bam files were analyzed by samtools. The number of reads aligned for each chromosome were calculated using samtools view. For further analysis, aligned reads were separated into two groups: mapped to the chrM or aligned to nuclear DNA. Mitochondrial reads were filtered from nuclear DNA of mitochondrial origin (nuMTs) by RtN (https://github.com/Ahhgust/RtN). For mtDNA content score calculation, we used the ratio (in percent) of read amounts aligned on chrM (mtDNA) to read amounts aligned on chr1-22 (autosomes). The complete sample inventory is openly available on Github (https://github.com/mtdnaPGT/PGT/blob/master/Embyo_data.xlsx). For mtDNA variant calling, vcf files were created by gatk tools (version 4.2.6.1) using AddOrReplaceReadGroups, MarkDuplicates, BaseRecalibrator, ApplyBQSR, Mutec2, LearnReadOrientationModel, and FilterMutectCalls algorithms. Mutec2 and FilterMutectCalls ran with parameters characteristic for mitochondrial genome analysis. For statistical analysis and its visualization, we used custom R-scripts (Mann-Whitney U test; glm; Wilcox test; Spearman correlation, point biserial correlation). All scripts and all data corresponding to mtDNA variant analysis are openly available on Github (https://github.com/mtdnaPGT/PGT).

