## Supplementary Data for "Aneuploidy in human embryos is associated with a maternal age-independent increase in mitochondrial DNA content and an enrichment of ultra-rare mitochondrial DNA variants"

### Supplementary Data 1

**The mtDNA content was found to be increased in NRFT compared to RFT embryos in independent analyses using both the VeriSeq PGS and AB-PGT protocols.**

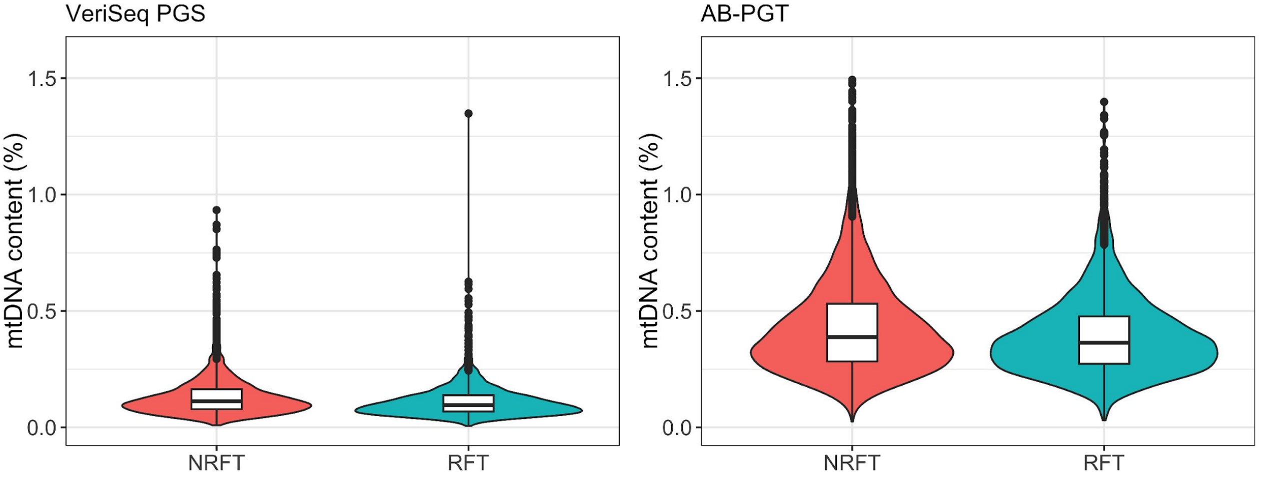


| Protocol | NRFT | | | RFT | | | Mann-Whitney U test | | Corrected *P*-value |
|---|---|---|---|---|---|---|---|---|---|
| | Sample size | Median | SD | Sample size | Median | SD | W | *P*-value | |
| VeriSeq PGS | 2009 | 0.112 | 0.095 | 1648 | 0.096 | 0.0776 | 1908340 | 1.70e-15 | 0.0008 |
| AB-PGT | 4199 | 0.388 | 0.212 | 3754 | 0.363 | 0.1740 | 8598874 | 2.25e-12 | 0.0262 |

A large sample size can affect *P*-values, obtained by the Mann-Whitney U test (Gómez-de-Mariscal E. et.al., 2021). To address this issue, we used the pMoSS tool (https://github.com/BIIG-UC3M/pMoSS) to estimate representative sample sizes of 600 for VeriSeq PGS and 1,100 for AB-PGT. Then, we performed 10,000 random samplings for both protocols, comparing 600 random NRFT with 600 random RFT from VeriSeq PGS and comparing 1,100 random NRFT with 1,100 RFT from AB-PGT. In the VeriSeq analysis, we observed 8 *P*-values that were higher than 0.05, leading to a corrected *P*-value of 0.0008 (8/10000). In the AB-PGT analysis, we observed 262 *P*-values that were higher than 0.05, leading to a corrected p-value of 0.0262 (262/10000).

Gómez-de-Mariscal E, Guerrero V, Sneider A, Jayatilaka H, Phillip JM, Wirtz D, Muñoz-Barrutia A. Use of the p-values as a size-dependent function to address practical differences when analyzing large datasets. Sci Rep. 2021 Oct 22;11(1):20942. doi: 10.1038/s41598-021-00199-5. PMID: 34686696; PMCID: PMC8536742.

#### Supplementary Data 2

**The mtDNA content was found to be higher in NRFT compared to RFT embryos in three different mtDNA intervals: the minor arc (442-5,720 nt), the major arc (1-109 nt and 5,799-16,569 nt), and the origins of replication (OH: 110-441 nt and OL: 5,721-5,798 nt) in independent analyses using both the VeriSeq PGS and AB-PGT protocols.**

VeriSeq PGS

| | NRFT | | | RFT | | | Mann-Whitney U test | | |
|---|---|---|---|---|---|---|---|---|---|
| **Intervals** | **Sample size** | **Median** | **SD** | **Sample size** | **Median** | **SD** | **W** | ***P*-value** | **Corrected P-value** |
| minor arc | 2009 | 0.027 | 0.0231 | 1648 | 0.023 | 0.0179 | 1892868 | 7.63e-14 | 0.0021 |
| major arc | 2009 | 0.053 | 0.0475 | 1648 | 0.045 | 0.0397 | 1900113 | 1.33e-14 | 0.0014 |
| OH and OL | 2009 | 0.037 | 0.0323 | 1648 | 0.031 | 0.0261 | 1891330 | 1.11e-13 | 0.0029 |

AB-PGT

| | NRFT | | | RFT | | | Mann-Whitney U test | | |
|---|---|---|---|---|---|---|---|---|---|
| **Intervals** | **Sample size** | **Median** | **SD** | **Sample size** | **Median** | **SD** | **W** | ***P*-value** | **Corrected *P*-value** |
| minor arc | 4199 | 0.148 | 0.08120 | 3754 | 0.139 | 0.06790 | 8568282 | 1.83e-11 | 0.0021 |
| major arc | 4199 | 0.235 | 0.12900 | 3754 | 0.221 | 0.10600 | 8606921 | 1.28e-12 | 0.0014 |
| OH and OL | 4199 | 0.011 | 0.00704 | 3754 | 0.010 | 0.00587 | 8603492 | 1.48e-12 | 0.0029 |

The derivation of the corrected *P*-values is explained in Supplementary Data 1.

### Supplementary Data 3

#### NRFT versus RFT embryos are characterized by increased maternal age and increased mtDNA content.

Multiple logistic regression analysis was performed using the glm function in R, with the dependent variable coded as 1 for NRFT embryos and 0 for RFT embryos and three continuous independent variables: mtDNA content, maternal age and parental age. The independent variables were **scaled** to allow for direct comparison of coefficients. The results were quantitatively similar for both the VeriSeq PGS and AB-PGT protocols, showing that maternal age was the strongest factor associated with aneuploidy, followed by mtDNA content. Paternal age was not significantly associated with aneuploidy. An additional analysis, which tested for an interaction between maternal age and mtDNA content (data not shown), did not reveal any interaction between these two variables (see also Supplementary Data 4). This indicates that mtDNA content is associated with aneuploidy independently of maternal age.

VeriSeq PGS

| Predictors | Odds Ratios (confidence intervals, CI) | Coefficients | *P*-value |
|---|---|---|---|
| (Intercept) | 1.33 (1.21-1.45) | 6.20 | <0.001 |
| Maternal age | 1.50 (1.35-1.66) | 7.68 | <0.001 |
| mtDNA content | 1.40 (1.26-1.57) | 5.97 | <0.001 |
| Paternal age | 1.05 (0.95-1.16) | 0.89 | 0.375 |

AB-PGT

| Predictors | Odds Ratios (confidence intervals,CI) | Coefficients | *P*-value |
|---|---|---|---|
| (Intercept) | 1.10 (1.05-1.16) | 3.78 | <0.001 |
| Maternal age | 1.52 (1.43-1.62) | 13.52 | <0.001 |
| mtDNA content | 1.23 (1.17-1.30) | 7.49 | <0.001 |
| Paternal age | 0.99 (0.93-1.05) | -0.45 | 0.650 |

#### Supplementary Data 4

**Aneuploidy is more strongly associated with maternal age and less strongly associated with mtDNA content, as indicated by point biserial correlations. Maternal age and mtDNA content are independent of each other, as shown by Spearman correlations. Similar results were also obtained through a logistic regression approach (Supplementary Data 3).**

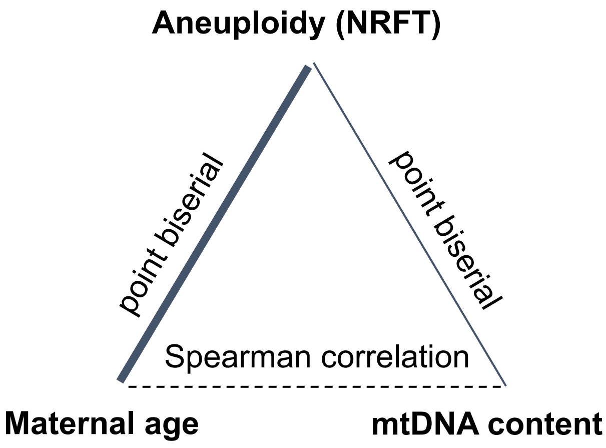


According to point biserial correlation (NRFT embryos were coded as 1 and NRFT embryos were coded as 0), both maternal age and mtDNA content are associated with aneuploidy. The strength of the association is 2 times stronger for maternal age compared to mtDNA content (see also Supplementary Data 3).

VeriSeq PGS

| | point biserial | 95% CI |
|---|---|---|
| Maternal age | 0.25 | [0.22, 0.28] |
| mtDNA content | 0.12 | [0.08, 0.15] |

AB-PGT

| | point biserial | 95% CI |
|---|---|---|
| Maternal age | 0.21 | [0.19, 0.23] |
| mtDNA content | 0.1 | [0.07, 0.12] |

According to Spearman rank correlations mtDNA content is independent of maternal age in all subsets of embryos and protocols (see also Supplementary Data 3) - especially when considering only the RFT or NRFT groups.

| Protocol (NRFT or/ans RFT) | Sample size | Spearman's Rho (*P*-value) |
|---|---|---|
| AB-PGT (NRFT, RFT) and VeriSeq PGS (NRFT, RFT) | 11351 | 0.061 (7.56e-11) |
| VeriSeq PGS (NRFT, RFT) | 3554 | 0.047 (0.005) |
| VeriSeq PGS (NRFT) | 1957 | 0.021 (0.39) |
| VeriSeq PGS (RFT) | 1597 | 0.012 (0.53) |
| AB-PGT (NRFT, RFT) | 7797 | 0.039 (0.0004) |
| AB-PGT (NRFT) | 4117 | 0.019 (0.22) |
| AB-PGT (RFT) | 3680 | 0.025 (0.12) |

#### Supplementary Data 5

**The risk of aneuploidy increases with both mtDNA content and maternal age (see statistical details in Supplementary Data 3 and 4). NRFT and RFT embryos are marked as red and blue points correspondingly.**

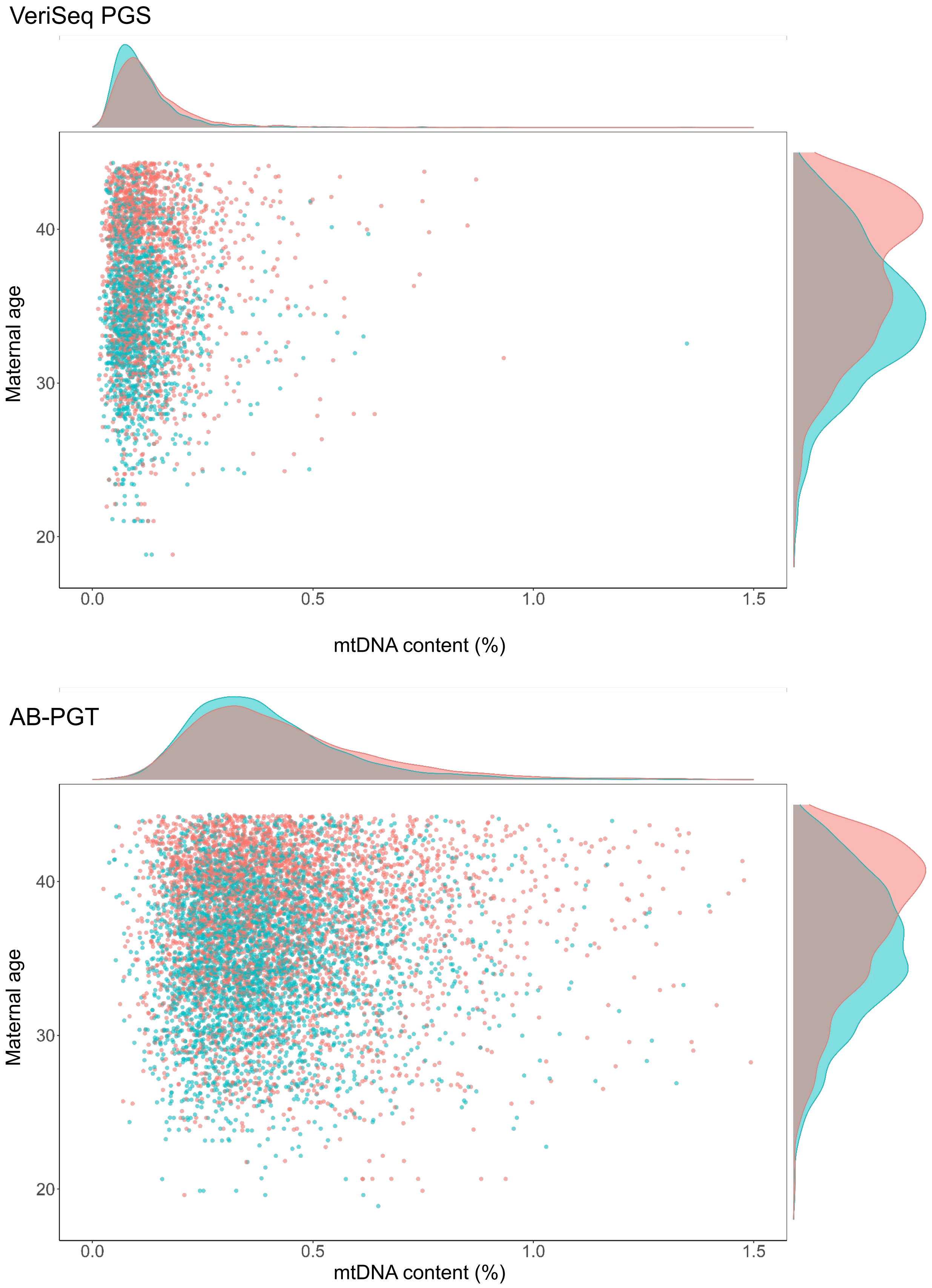

**Supplementary Data 6**

The fraction of NRFT embryos increases with both mtDNA content and maternal age, as shown in the cumulative frequency graphs. The y-axis displays the fraction of NRFT embryos (%), corresponding to all samples with an X value equal to or greater than the current X value (either mtDNA content or maternal age).

The x-axis reflects the percentile of mtDNA content (%). Among embryos with the highest mtDNA content (for example, the 95th percentile), we observed 89% (for VeriSeq PGS) and 67% (for AB-PGT) to be NRFT embryos.

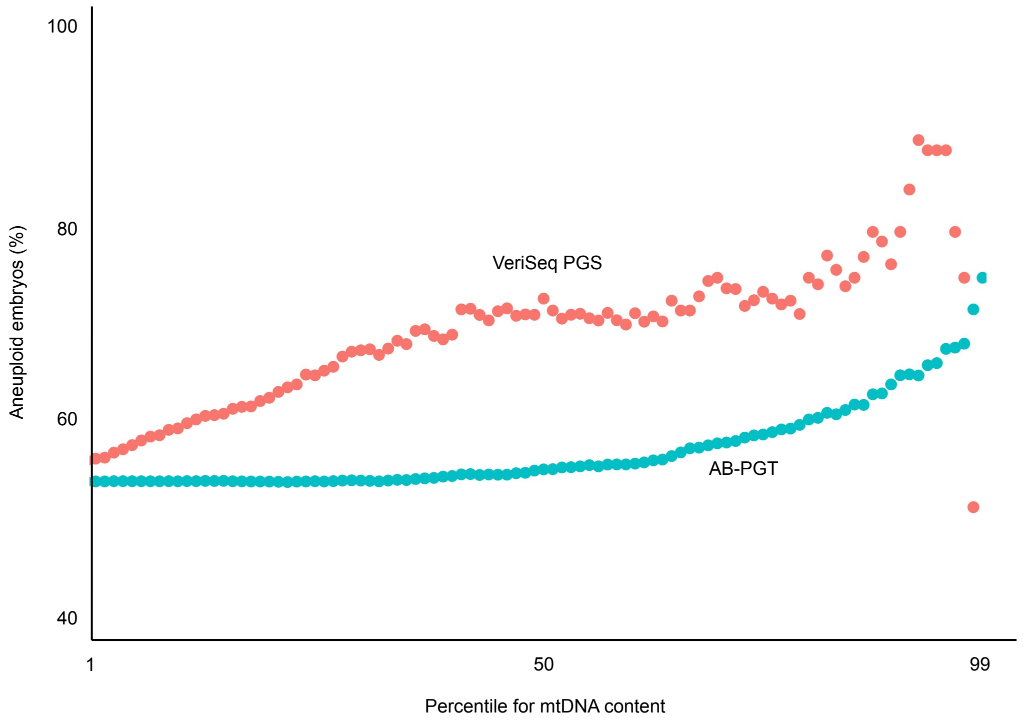

#### Supplementary Data 6 (additional 1)

The x-axis shows maternal age (for both VeriSeq PGS and AB-PGT). As maternal age increases, the fraction of NRFT embryos also increases.

The value of each points are presented in the table below.

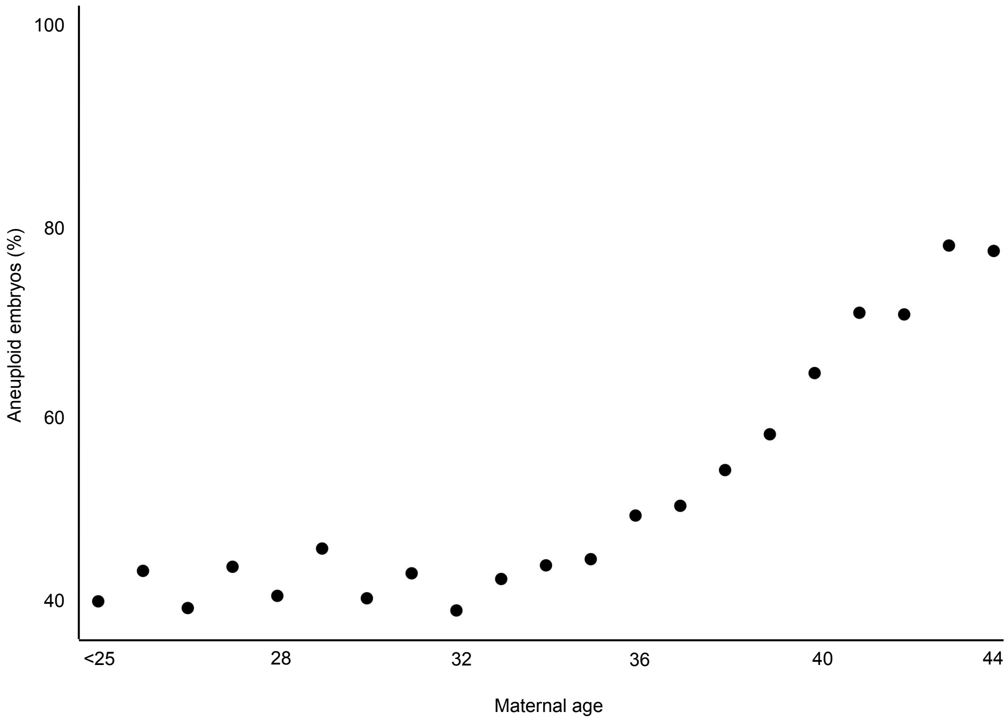


| Maternal age | < 25 | 25 | 26 | 27 | 28 | 29 | 30 | 31 | 32 | 33 | 34 | 35 | 36 | 37 | 38 | 39 | 40 | 41 | 42 | 43 | 44 |
|---|---|---|---|---|---|---|---|---|---|---|---|---|---|---|---|---|---|---|---|---|---|
| **Number of NRFT embryos** | 50 | 36 | 46 | 87 | 95 | 136 | 166 | 167 | 220 | 305 | 324 | 346 | 388 | 409 | 414 | 452 | 541 | 587 | 494 | 459 | 352 |
| **Number of RFT embryos** | 76 | 48 | 72 | 114 | 141 | 165 | 249 | 225 | 348 | 421 | 424 | 437 | 410 | 415 | 361 | 339 | 312 | 255 | 296 | 139 | 110 |
| **(NRFT/(RFT+NRFT))*100** | 40 | 43 | 39 | 43 | 40 | 45 | 40 | 43 | 39 | 42 | 43 | 44 | 49 | 50 | 53 | 57 | 63 | 70 | 70 | 77 | 76 |

#### Supplementary Data 7

**A 10-year increase in maternal age is equivalent to an increase of 0.6% in mtDNA content in terms of the risk of aneuploidy.**

Both maternal age and mtDNA content are associated with an increased risk of aneuploidies. To estimate quantitatively the equivalent changes in maternal age and mtDNA content, assuming their linear effects on aneuploidy, we used estimated coefficients from multiple logistic regression where NRFT (coded as one) was a function of two unscaled variables: maternal age and mtDNA content. Obtained odds ratios (see table below), were used to calculate ΔmtDNA content associated with Δmaternal age (see formula below).

VeriSeq PGS

| Predictors | Odds Ratios (confidence intervals, CI) | Coefficients | *P*-value |
|---|---|---|---|
| (Intercept) | 0.02 (0.01 – 0.33) | -14.54 | <0.001 |
| Maternal age | 1.12 (1.10 – 1.13) | 14.40 | <0.001 |
| mtDNA content | 18.67 (7.76 – 46.30) | 6.42 | <0.001 |

AB- PGT

| Predictors | Odds Ratios (confidence intervals, CI) | Coefficients | *P*-value |
|---|---|---|---|
| (Intercept) | 0.03 (0.02 – 0.04) | -18.80 | <0.001 |
| Maternal age | 1.09 (1.08 – 1.10) | 17.96 | <0.001 |
| mtDNA content | 2.83 (2.22 – 3.61) | 8.38 | <0.001 |

$$\Delta\text{maternal age} = \Delta\text{mtDNA content} \cdot \frac{\ln\text{ (Odds Ratios of mtDNA content)}}{\ln\text{ (Odds Ratios of Maternal age)}}$$

VeriSeq PGS
Δmaternal age = 10 years (for example)
ln (Odds Ratios of mtDNA content) = ln(1.12) = 0.11
ln (Odds Ratios of Maternal age) = ln(18.67) = 2.93
ΔmtDNA content = 0.376%

AB-PGT
Δmaternal age = 10 years (for example)
ln (Odds Ratios of mtDNA content) = ln(1.09) = 0.086
ln (Odds Ratios of Maternal age) = ln(2.83) = 1.04
ΔmtDNA content = 0.827%

In Figure 4 of the main text, we used the same logic to draw a purple arrow indicating the equivalent changes. Specifically, for VeriSeq PGS, every 5-year increase in maternal age is equivalent to an increase of 0.188% in mtDNA content.

### Supplementary Data 8

**The increased mtDNA content in NRFT embryos is primarily due to an increase in the number of reads mapped to the mtDNA chromosome (hereafter “chrM coverage”), rather than a decrease in the number of reads mapped to the autosomes (hereafter “chr1-22 coverage”)**

The mtDNA content, which is defined as a ratio of the chrM coverage to the chr1-22 coverage, is affected by both these numbers. According to our biological interpretation, the increased mtDNA content in NRFT embryos is primarily due to an increase in chrM coverage, rather than a decrease in chr1-22 coverage. To test this, we performed several analyses.

A) chrM coverage is higher in NRFT versus RFT embryos, while chr1-22 coverage show no difference: mean

VeriSeq PGS

| | | Number of reads | | |
|---|---|---|---|---|
| **Type** | **Samples size** | **Mean (chr1-22)** | **Mean (chrM)** | **Mean (chrM) / Mean (chr1-22)** |
| NRFT | 2009 | 702226 | 950 | 0.0014 |
| RFT | 1648 | 697628 | 797 | 0.0011 |

AB-PGT

| | | Number of reads | | |
|---|---|---|---|---|
| **Type** | **Samples size** | **Mean (chr1-22)** | **Mean (chrM)** | **Mean (chrM) / Mean (chr1-22)** |
| NRFT | 4199 | 288848 | 1237 | 0.0043 |
| RFT | 3754 | 291411 | 1138 | 0.0039 |

B) chrM coverage is higher in NRFT versus RFT embryos, while chr1-22 coverage show no difference: p-values derived from Mann-Whitney U test between NRFT and RFT groups.

| | P-value | |
|---|---|---|
| **Protocol** | **chr1-22** | **chrM** |
| VeriSeq PGS | 0.46520 | 2.052e-15 |
| AB-PGT | 0.08447 | 2.217e-09 |

C) mtDNA content correlates stronger with chrM coverage than with the chr1-22 coverage (Spearman’s rank correlations)

| | VeriSeq PGS | | AB-PGT | |
|---|---|---|---|---|
| **Comparisons** | **rho** | ***P*-value** | **rho** | ***P*-value** |
| chrM VS. chr1-22 | 0.19 | < 2.2e-16 | 0.39 | < 2.2e-16 |
| chrM VS. mtDNA content | 0.97 | < 2.2e-16 | 0.87 | < 2.2e-16 |
| chr1-22 VS. mtDNA content | -0.009 | 0.57 | -0.08 | 4.149e-12 |

### Supplementary Data 8 (additional 1)

**To ensure that the coverage of chr1-22 does not impact our results, we reran multiple logistic regression analyses (as described in Supplementary Data 3) with an additional factor, the coverage of chr1-22. For both platforms, this new factor was not significant. As in the Supplementary Data 3, NRFT embryos were coded as one and RFT embryos were coded as zero.**

#### VeriSeq PGS

| Predictors | Odds Ratios (confidence intervals, CI) | Coefficients | *P*-value |
| --- | --- | --- | --- |
| (Intercept) | 1.25 (1.16-1.33) | 6.24 | <0.001 |
| Maternal age | 1.69 (1.58-1.82) | 14.38 | <0.001 |
| mtDNA content | 1.30 (1.20-1.40) | 6.43 | <0.001 |
| chr1-22 | 1.05 (0.98-1.13) | 1.32 | 0.186 |

#### AB-PGT

| Predictors | Odds Ratios (confidence intervals, CI) | Coefficients | *P*-value |
| --- | --- | --- | --- |
| (Intercept) | 1.12 (1.07-1.18) | 5.04 | <0.001 |
| Maternal age | 1.54 (1.47-1.61) | 17.93 | <0.001 |
| mtDNA content | 1.22 (1.17-1.28) | 8.28 | <0.001 |
| chr1-22 | 0.98 (0.94-1.03) | -0.79 | 0.431 |

### Supplementary Data 9

**The mtDNA content is consistently higher in NRFT embryos compared to RFT embryos irrespective of additional factors, such as the stage of embryo development, molecular karyotype, morphological groups, and type of chromosomal abnormality.**

Certain additional traits of embryos, such as the stage of development, may be correlated with mtDNA content and thus influence our observations. To control for these potential biases, we employed two approaches. First, we divided all embryos into groups and demonstrated that the increase in mtDNA content of NRFT versus RFT embryos remains consistent within each group (Supplementary Data 9). Additionally, we conducted multiple logistic regression analysis while controlling for these extra factors (Supplementary Data 10). These efforts helped ensure the robustness of our findings.

A) Embryo development day (5, 6, 7). The mtDNA content decreases over the course of embryo development from day 5 to day 7. However, within each day, the mtDNA content is consistently higher in NFRT embryos compared to RFT embryos.

VeriSeq PGS

| | NRFT | | | RFT | | | Mann-Whitney U test | |
|---|---|---|---|---|---|---|---|---|
| **Embryo development day** | **Sample size** | **Median** | **SD** | **Sample size** | **Median** | **SD** | **W** | ***P*-value** |
| 5 | 1345 | 0.125 | 0.1010 | 1230 | 0.107 | 0.0828 | 964876.5 | 2.73e-13 |
| 6 | 623 | 0.089 | 0.0702 | 393 | 0.069 | 0.0449 | 152366.5 | 4.89e-11 |
| 7 | 17 | 0.089 | 0.0496 | 14 | 0.051 | 0.0166 | 217 | 1.08e-04 |

AB-PGT

| | NRFT | | | RFT | | | Mann-Whitney U test | |
|---|---|---|---|---|---|---|---|---|
| **Embryo development day** | **Sample size** | **Median** | **SD** | **Sample size** | **Median** | **SD** | **W** | ***P*-value** |
| 5 | 2690 | 0.419 | 0.223 | 2684 | 0.388 | 0.184 | 3952497 | 1.71e-09 |
| 6 | 1411 | 0.339 | 0.178 | 1013 | 0.306 | 0.122 | 837488 | 4.96e-13 |
| 7 | 71 | 0.300 | 0.123 | 34 | 0.272 | 0.129 | 1363.5 | 2.85e-01 |

### Supplementary Data 9 (additional 1)

B) Embryo molecular karyotype (XX and XY). For each molecular karyotype the mtDNA content is consistently higher in NFRT embryos compared to RFT embryos.

#### VeriSeq PGS

| | NRFT | | | RFT | | | Mann-Whitney U test | |
|---|---|---|---|---|---|---|---|---|
| **Embryo molecular karyotype** | **Sample size** | **Median** | **SD** | **Sample size** | **Median** | **SD** | **W** | ***P*-value** |
| XX | 925 | 0.115 | 0.0938 | 795 | 0.094 | 0.0875 | 439449 | 2.79e-12 |
| XY | 1031 | 0.110 | 0.0911 | 853 | 0.098 | 0.0671 | 486504 | 6.88e-05 |

#### AB-PGT

| | NRFT | | | RFT | | | Mann-Whitney U test | |
|---|---|---|---|---|---|---|---|---|
| **Embryo molecular karyotype** | **Sample size** | **Median** | **SD** | **Sample size** | **Median** | **SD** | **W** | ***P*-value** |
| XX | 1923 | 0.388 | 0.216 | 1852 | 0.363 | 0.182 | 1935515 | 3.75e-06 |
| XY | 2187 | 0.389 | 0.209 | 1902 | 0.365 | 0.166 | 2288554 | 2.97e-08 |

### Supplementary Data 9 (additional 2)

C) Morphological groups. Samples were divided into five groups based on Gardner morphological classification. This system assigns three separate quality scores to each blastocyst embryo: expansion grade, inner cell mass (ICM) grade, and trophectoderm (TE) grade. For each of the five groups, the mtDNA content is consistently higher in NFRT embryos compared to RFT embryos

| Expansion grade | Blastocyst development and stage status |
|---|---|
| 1 | Blastocoel cavity less than half the volume of the embryo |
| 2 | Blastocoel cavity more than half the volume of the embryo |
| 3 | Full blastocyst, cavity completely filling the embryo |
| 4 | Expanded blastocyst, cavity larger than the embryo, with thinning of the shell |
| 5 | Hatching out of the shell |
| 6 | Hatched out of the shell |

**Example**

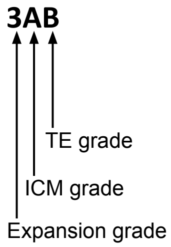


| ICM grade | Inner cell mass quality |
|---|---|
| A | Many cells, tightly packed |
| B | Several cells, loosely grouped |
| C | Very few cells |

| TE grade | Trophectoderm quality |
|---|---|
| A | Many cells, forming a cohesive layer |
| B | Few cells, forming a loose epithelium |
| C | Very few large cells |

#### VeriSeq PGS

| | NRFT | | | RFT | | | Mann-Whitney U test | |
|---|---|---|---|---|---|---|---|---|
| Groups | Sample size | Median | SD | Sample sizes | Median | SD | W | *P*-value |
| (3-6)AA | 415 | 0.113 | 0.0862 | 545 | 0.100 | 0.0679 | 129286.0 | 1.41e-04 |
| (3-6)AA and (3-6)AB and (3-6)BA * | 981 | 0.114 | 0.0925 | 1063 | 0.100 | 0.0712 | 607630.0 | 9.90e-11 |
| (3-6)AB and (3-6)BA | 566 | 0.116 | 0.0967 | 518 | 0.099 | 0.0746 | 173300.5 | 2.14e-07 |
| (3-6)BB | 561 | 0.104 | 0.0916 | 378 | 0.086 | 0.0647 | 127785.5 | 9.39e-08 |
| (3-6)BC and (3-6)CB and (3-6)CC | 116 | 0.102 | 0.0750 | 56 | 0.070 | 0.0778 | 4075.5 | 6.88e-03 |

#### AB-PGT

| | NRFT | | | RFT | | | Mann-Whitney U test | |
|---|---|---|---|---|---|---|---|---|
| Groups | Sample size | Median | SD | Sample sizes | Median | SD | W | *P*-value |
| (3-6)AA | 905 | 0.393 | 0.205 | 1309 | 0.372 | 0.181 | 637820.5 | 2.09e-03 |
| (3-6)AA and (3-6)AB and (3-6)BA * | 2266 | 0.397 | 0.212 | 2516 | 0.375 | 0.177 | 3069782.0 | 4.28e-06 |
| (3-6)AB and (3-6)BA | 1361 | 0.399 | 0.216 | 1207 | 0.378 | 0.173 | 881025.5 | 1.47e-03 |
| (3-6)BB | 1448 | 0.381 | 0.211 | 990 | 0.336 | 0.166 | 831730.5 | 1.63e-11 |
| (3-6)BC and (3-6)CB and (3-6)CC | 197 | 0.366 | 0.204 | 87 | 0.315 | 0.162 | 9867.0 | 4.21e-02 |

* this group combines two other groups: "(3-6)AA" and "(3-6)AB and (3-6)BA"

### Supplementary Data 9 (additional 3)

D) Chromosomal abnormality. For both protocols mtDNA content is higher in NRFT versus RFT embryos irrespective of the type of chromosomal abnormality, observed in NRFT embryos: insertions (ADD), deletions (LOST) and mosaic type samples (MOSAIC).

#### VeriSeq PGS

| | NRFT | | | RFT | | | Mann-Whitney U test | |
|---|---|---|---|---|---|---|---|---|
| **Comparison** | **Sample size** | **Median** | **SD** | **Sample size** | **Median** | **SD** | **W** | ***P*-value** |
| RFT VS. ADD | 467 | 0.102 | 0.0929 | 1648 | 0.096 | 0.0776 | 411049 | 2.43e-02 |
| RFT VS. LOST | 873 | 0.117 | 0.0984 | 1648 | 0.096 | 0.0776 | 867529 | 1.57e-17 |
| RFT VS. MOSAIC | 558 | 0.112 | 0.0877 | 1648 | 0.096 | 0.0776 | 527591 | 1.85e-07 |

#### AB-PGT

| | NRFT | | | RFT | | | Mann-Whitney U test | |
|---|---|---|---|---|---|---|---|---|
| **Comparison** | **Sample size** | **Median** | **SD** | **Sample size** | **Median** | **SD** | **W** | ***P*-value** |
| RFT VS. ADD | 1122 | 0.371 | 0.2 | 3754 | 0.363 | 0.174 | 2188190 | 4.7e-02 |
| RFT VS. LOST | 2101 | 0.399 | 0.218 | 3754 | 0.363 | 0.174 | 4474002 | 1.23e-17 |
| RFT VS. MOSAIC | 894 | 0.387 | 0.209 | 3754 | 0.363 | 0.174 | 1802819 | 5.39e-04 |

#### Supplementary Data 10

To ensure additionally that extra factors (morphology, molecular karyotype, development day, protocol) don't impact our results, we reran multiple logistic regression analyses (as described in Supplementary Data 3) including:

- morphology group (“AA” versus all others);
- molecular karyotype (XX versus XY);
- embryo development day: early days (4 and 5) versus late days (6 and 7);
- protocol (AB-PGT versus VeriSeq PGS).

Irrespective of all these extra factors, added one by one or altogether, multiple logistic regression models demonstrated always that maternal age is the first strongest factor and mtDNA content is the second strongest one, associated with NRFT embryos.

#### Supplementary Data 11

**Family-based analyses show that NRFT embryos have a higher mtDNA content compared to RFT embryos within the same siblings.**

We compared the **mean** mtDNA content between NRFT and RFT embryos in 1,827 families with at least one NRFT and at least one RFT embryo. The results of the paired Mann-Whitney U test showed that the null hypothesis of no difference in mtDNA content between NRFT and RFT embryos was rejected, indicating that NRFT embryos have a higher mtDNA content than RFT embryos.

| | | NRFT VS. RFT | | | | NRFT / RFT | |
|---|---|---|---|---|---|---|---|
| | | paired Mann-Whitney U test | | Wilcox test (mu=1) | | | |
| Protocol | Pairs | V | P-value | V | P-value | Median | Mean |
| VeriSeq PGS | 605 | 122143.5 | 1.94e-13 | 132218.0 | 1.33e-22 | 1.189 | 1.396 |
| AB-PGT | 1222 | 444782.0 | 4.10e-09 | 474536.5 | 5.32e-17 | 1.067 | 1.211 |
| VeriSeq PGS and AB-PGT* | 1827 | 1014171.0 | 2.77e-16 | 1111214.0 | 3.56e-36 | 1.095 | 1.272 |

Out of 1,827 families, only 1,784 have information on maternal age. These 1,784 families were divided into two groups based on whether the maternal age was above or below the median (37 years). The results for these two groups were similar to the results for the full sample of 1,827 families.

The subset of families with maternal age greater than or equal to 37 years:

| | | NRFT VS. RFT | | | | NRFT / RFT | |
|---|---|---|---|---|---|---|---|
| | | paired Mann-Whitney U test | | Wilcox test (mu=1) | | | |
| Protocol | Pairs | V | p-value | V | p-value | Median | Mean |
| VeriSeq PGS | 243 | 19436.5 | 1.40e-05 | 20997.0 | 3.86e-09 | 1.172 | 1.387 |
| AB-PGT | 597 | 107915.5 | 9.57e-06 | 114937.5 | 5.57e-10 | 1.082 | 1.214 |
| VeriSeq PGS and AB-PGT* | 840 | 215622.0 | 1.96e-08 | 234393.5 | 5.71e-17 | 1.098 | 1.264 |

The subset of families with maternal age less than 37 years:

| | | NRFT VS. RFT | | | | NRFT / RFT | |
|---|---|---|---|---|---|---|---|
| | | paired Mann-Whitney U test | | Wilcox test (mu=1) | | | |
| Protocol | Pairs | V | p-value | V | p-value | Median | Mean |
| VeriSeq PGS | 342 | 38793.5 | 1.22e-07 | 41967 | 1.02e-12 | 1.175 | 1.369 |
| AB-PGT | 602 | 106079.5 | 1.77e-04 | 113284 | 2.58e-08 | 1.063 | 1.203 |
| VeriSeq PGS and AB-PGT* | 944 | 267750.0 | 3.14e-08 | 293944 | 2.08e-18 | 1.085 | 1.263 |

* In the case of paired Mann-Whitney U test, it is appropriate to merge familial data that has been analyzed using both protocols because each comparison pair belongs to the same protocol.

#### Supplementary Data 11 (additional 1)

**There is an increased mtDNA content in NRFT versus RFT embryos within each family irrespective of the maternal age.**

Each of the 1,784 families is represented by a line connecting the mean mtDNA content of NRFT and the mean mtDNA content of RFT embryos. The top figure shows 1,025 families (664 from AB, 361 from VeriSeq) where the mtDNA content of NRFT is greater than that of RFT embryos. The bottom figure shows 759 families (535 from AB-PGT, 224 from VeriSeq PGS) where the mtDNA content of NRFT is less than that of RFT embryos. The figure at the next page shows all 1,784 families, represented by the ratio of mtDNA content of NRFT to RFT embryos. By visual inspection, we can see that there is an excess of families with a ratio greater than 0, and this excess is uniformly distributed across different maternal ages (see next page).

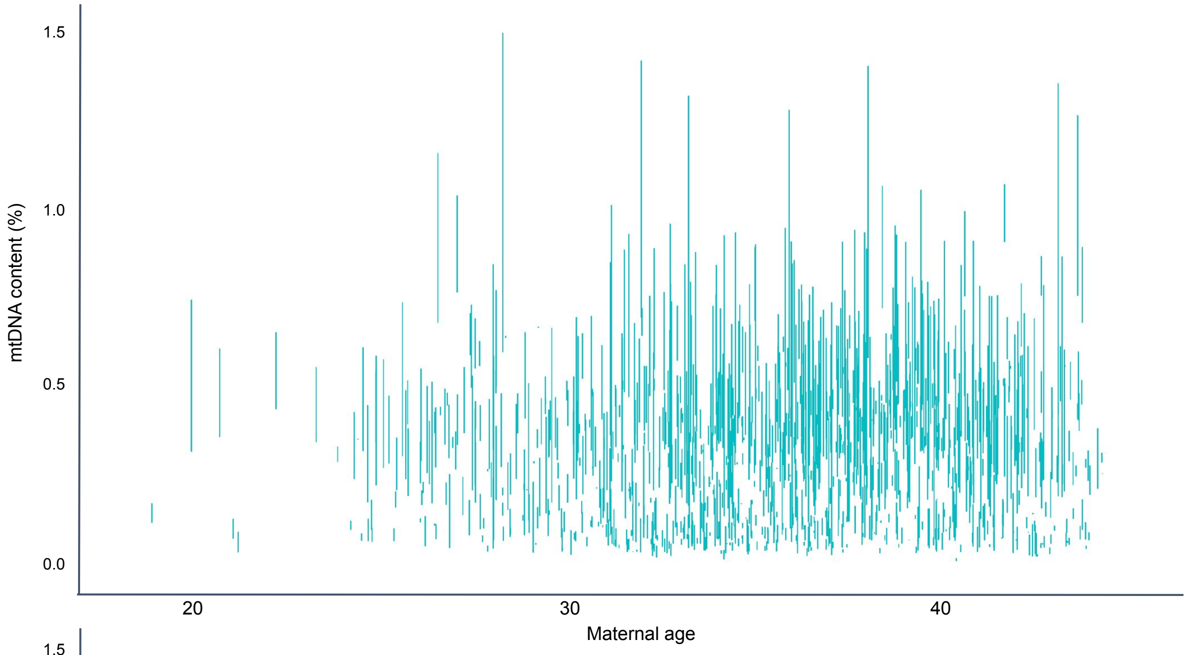


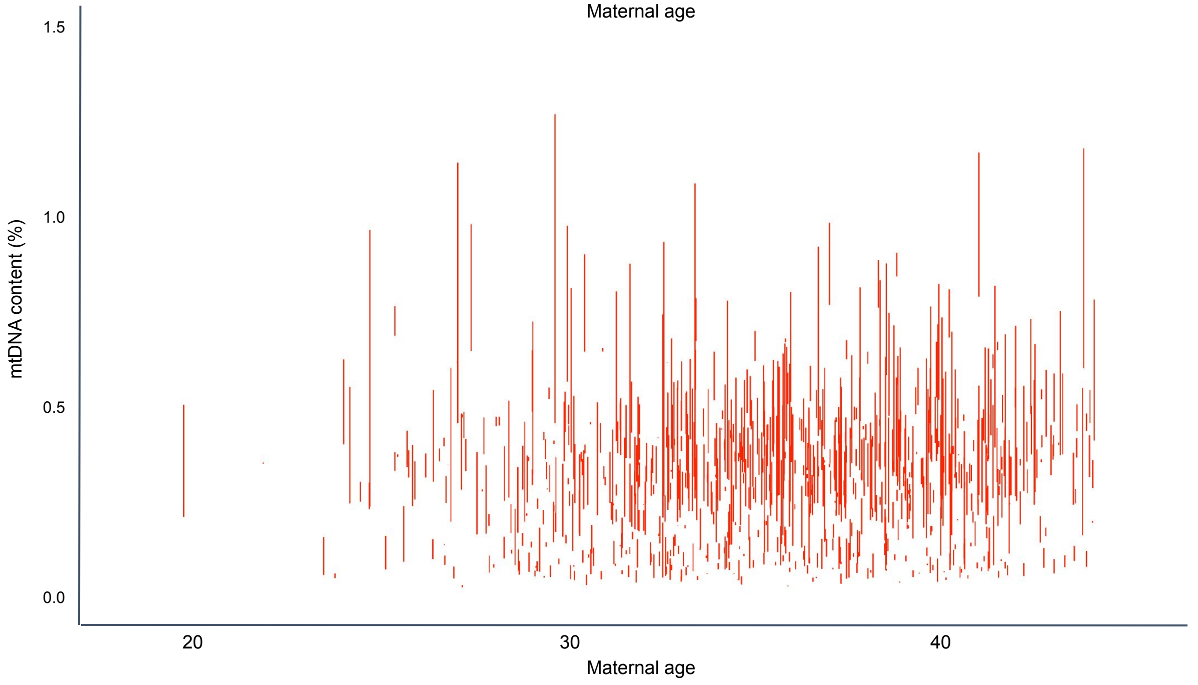

### Supplementary Data 11 (additional 2)

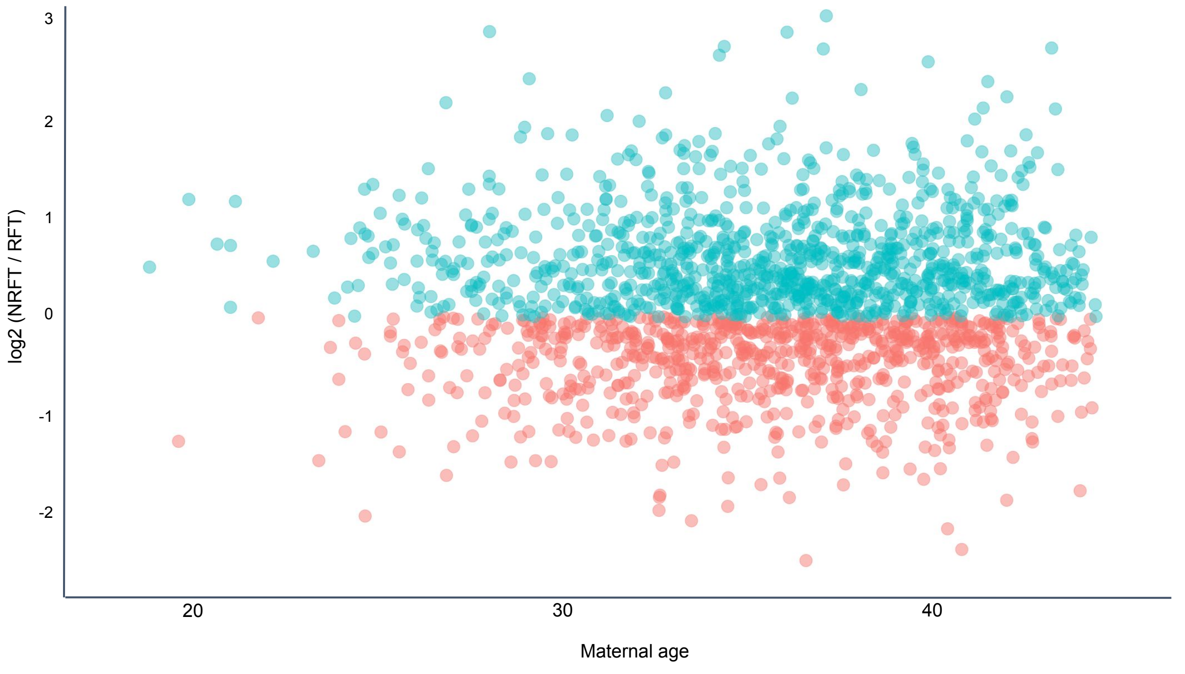

#### Supplementary Data 12

**The mitochondrial DNA of NRFT embryos is enriched in ultra-rare potentially deleterious variants: population-based analysis.**

We analyzed 11,610 vcf files of chrM, resulting in a total of 912,100 rows of concatenated data. The median coverage for called alternative variants was 2 for AB-PGT and 5 for VeriSeq PGS. After keeping only alternative mtDNA variants that were supported by at least one forward and one backward read, we were left with 44,912 rows.

In order to analyze the distribution of potentially-deleterious variants among NRFT and RFT embryos, we classified all alternative variants into three categories based on the frequencies of observed polymorphisms in the human population using 195,983 fully sequenced mitochondrial genomes from around the world (HELIX database, https://www.helix.com/pages/mitochondrial-variant-database, version 2020/03/27). The categories were:

- often altered positions (with three or more carriers of the alternative allele at a given position);
- rarely altered positions (with at least one carrier of the heteroplasmic alternative variant and less than three carriers of the homoplasmic variant);
- never altered positions (positions without any alternative variants, 6,316 positions).

We then performed a logistic regression by analyzing polymorphic sites (coded as 1 and 0) as a function of the embryo status (NRFT embryos were coded as 1 and RFT as 0). As a result of three logistic regression we found that only never altered positions were enriched in NRFT embryos (as shown by the positive 'Estimate' and significant p-value).

*often altered positions*

| Variable | Coefficients | Std. Error | z value | Pr(>\|z\|) |
|---|---|---|---|---|
| Intercept | 0.506651 | 0.003542 | 143.032 | <2e-16*** |
| NRFT | -0.001788 | 0.004749 | -0.377 | 0.706 |

*rarely altered positions*

| Variable | Coefficients | Std. Error | z value | Pr(>\|z\|) |
|---|---|---|---|---|
| Intercept | 0.416755 | 0.003489 | 119.465 | <2e-16*** |
| NRFT | -0.006494 | 0.004677 | -1.389 | 0.165 |

*never altered positions*

| Variable | Coefficients | Std. Error | z value | Pr(>\|z\|) |
|---|---|---|---|---|
| Intercept | 0.076595 | 0.001935 | 39.584 | <2e-16*** |
| NRFT | 0.008282 | 0.002594 | 3.193 | 0.00141** |

### Supplementary Data 12 (additional 1)

**The mitochondrial DNA of NRFT embryos is enriched in ultra-rare potentially deleterious variants: population-based analysis.**

Next, we delved deeper into the category of alternative variants located in "never altered positions". We described this category as a function of several independent variables:

- the embryo status (NRFT embryos were coded as 1 and RFT as 0);
- the total mtDNA coverage at a given site (continuous variable);
- the platform (VeriSeq PGS was coded as 1 and AB-PGT as 0).

Despite the fact that both the additional independent variables "total coverage in a site" and "platform" were significantly associated with never altered positions, we still observed a positive and significant association with NRFT. Therefore, we can see that both total coverage and platform did not eliminate the signal of the never altered positions.

| Variable | Coefficients | Std. Error | z value | Pr(>\|z\|) |
|---|---|---|---|---|
| Intercept | 0.069649 | 0.003154 | 22.08 | <2e-16*** |
| NRFT | 0.005814 | 0.002546 | 2.283 | 0.0224* |
| VeriSeq PGS | 0.134800 | 0.003383 | 39.848 | <2e-16*** |
| Coverage | -0.023028 | 0.001498 | -15.375 | <2e-16*** |

Next, we focused on the VeriSeq platform (19,623 rows out of the 44,912 used in previous analyses) and compared the recurrency of alternative alleles (confirmed by at least one forward and at least one reverse read) in NRFT and RFT embryos. The VeriSeq platform was chosen because it is characterized by highly covered "islands" within which alternative alleles can be called with high quality.

The 19,623 rows used in this analysis were split into 11,296 in NRFT and 8,327 in RFT embryos. Next, we concentrated on unique positions, resulting in 1,090 and 882 respectively. The overlap of these two lists gave us a total of 1,371 unique positions. For each of these positions, we counted the number of carriers of the alternative variant among NRFT and RFT embryos. Based on these two numbers, we determined a potential deleterious rank for each variant as described below.

We sorted the list of sites by:
(i) the descending ratio of the number of NRFT embryos carriers to the number of RFT ones;
(ii) by the descending number of NRFT embryo carriers;
(iii) by the ascending number of RFT embryo carriers.

Finally, we ranked all of these sites so that a high rank indicates an excess of the altered position among NRFT versus RFT embryos. The final table (n = 1,371) had a structure as shown below, with 618 often altered positions, 451 rarely altered positions, and 302 never altered positions.

### Supplementary Data 12 (additional 2)

| Position in mtDNA | Number of carriers among NRFT | Number of carriers among RFT | Number of carriers among NRFT / Number of carriers among RFT | Never altered position | Rarely altered position | Often altered position | Deleterious rank |
|---|---|---|---|---|---|---|---|
| 129 | 12 | 0 | Inf | 1 | 0 | 0 | 1371 |
| 214 | 10 | 0 | Inf | 0 | 0 | 1 | 1370 |
| 252 | 7 | 0 | Inf | 0 | 0 | 1 | 1369 |
| … | … | … | … | … | … | … | … |
| 16274 | 7 | 0 | Inf | 0 | 0 | 1 | 1368 |
| 494 | 4 | 4 | 1 | 0 | 0 | 1 | 518 |
| 634 | 4 | 4 | 1 | 0 | 1 | 0 | 517 |
| 3337 | 4 | 4 | 1 | 0 | 0 | 1 | 516 |
| 9983 | 4 | 4 | 1 | 0 | 0 | 1 | 515 |
| … | … | … | … | … | … | … | … |
| 745 | 0 | 5 | 0 | 0 | 1 | 0 | 4 |
| 10083 | 0 | 5 | 0 | 0 | 0 | 1 | 3 |
| 10256 | 0 | 5 | 0 | 0 | 0 | 1 | 2 |
| 3429 | 0 | 6 | 0 | 0 | 0 | 1 | 1 |

To analyze the potentially deleterious category of 'never altered' positions as a function of the ranked sites, we performed a logistic regression where never altered positions were coded as 1 and altered positions were coded as 0. We observed that never altered positions were positively associated with the rank (*P*-value = 0.007), meaning that among highly ranked sites (i.e., sites altered in NRFT but not in RFT), there is an excess of never altered positions. In other words, this is consistent with our hypothesis that mtDNA variants predominantly observed in NRFT (high rank) are enriched in deleterious variants (substitutions in never altered positions).

Next, we repeated the same analysis considering only SNPs (n = 1,165; 206 indels were deleted). The results were qualitatively similar (*P*-value = 0.00385).

Finally, we analyzed the distribution of control polymorphic categories (often and rarely altered) as a function of the deleterious rank. We did not observe any associations for the whole set of variants (n = 1,371) or for SNPs only (n = 1,165):
often altered positions: *P*-value = 0.284 (SNPs and Indels), *P*-value = 0.165 (SNPs);
rarely altered positions: *P*-value = 0.206 (SNPs and Indels), *P*-value = 0.319 (SNPs).

#### Supplementary Data 13

**The mitochondrial DNA of NRFT embryos is enriched in ultra-rare potentially deleterious variants: family-based analysis.**

11,610 bam files of chrM were analyzed to calculate two parameters for each file: the total coverage of mtDNA reads and the number of "mutated" mtDNA reads. A read was considered "mutated" if it (1) had an alternative allele on a position that had not previously been altered (according to the HELIX database; see Supplementary Data 12) and (2) this alternative allele was supported by at least one forward and one backward read.

We categorized the total number of mtDNA reads for each of 1,827 families with at least one NRFT and at least one RFT embryo into four categories: the number of "mutated" reads in NRFT, the number of all reads in NRFT, the number of "mutated" reads in RFT, and the number of all reads in RFT. Using those four numbers for each family we calculated the ratio of the total number of "mutated" mtDNA reads to the total number of all mtDNA reads for the NRFT and RFT subsets separately. Paired Mann-Whitney U test demonstrated significant excess of the fraction of mutated reads in NRFT versus RFT in the whole dataset (*P*-value = 0.0046, N = 1,827), however when we split dataset into two platforms only AB-PGT demonstrated the effect (*P*-value = 0.0007, n = 1,222), while VeriSeq PGS, probably due to low sample size, did not show the trend (*P*-value = 0.96, n = 605).

For the following tests, we eliminated families that contained zeroes: 141 out of the 1,827 families were excluded from the analysis because they had no mutated reads. 8 families had no "mutated" mtDNA reads for NRFT and RFT, 75 families had no "mutated" mtDNA reads for NRFT, and 58 families had no "mutated" mtDNA reads for RFT. For the remaining 1,686 families (1,827 minus 141), we plotted the ratios (the fraction of "mutated" mtDNA reads in NRFT versus RFT), on a histogram (see Fig. 3b). The median and mean of the ratio were significantly higher than the expected value one (*P*-value < 2.2e-16, according to a Wilcoxon test with mu equals 1).

We further analyzed 1,646 families based on maternal age, excluding 40 families out of the 1,686 due to missing information on maternal age. Visual inspection of the data (Fig. 3c) shows that there is a higher than expected number of families with a ratio greater than 0, and this excess is evenly distributed across different maternal ages. Out of the 1,646 families, 1,113 were analyzed using the AB-PGT platform and 533 were analyzed using the VeriSeq PGS platform. Only the families analyzed using the AB-PGT platform showed an excess of mutated reads in NRFT compared to RFT. Out of the 1,113 families analyzed using AB-PGT, an excess of mutated reads in NRFT compared to RFT was observed in 55% (613 families). Among the 533 families analyzed using VeriSeq PGS, 49.7% (265 families) showed an excess of mutated reads in NRFT compared to RFT. Two plots (top and middle) illustrating these results can be found on the following page.

We wanted to determine whether the excess of mutated reads in NRFT (i.e. log2(NRFT/RFT)>0) is dependent on the magnitude of the ratio (i.e. abs(log2(NRFT/RFT))). To do this, we plotted log2(NRFT/RFT) as a function of abs(log2(NRFT/RFT)). Our results showed that in families with a large contrast between NRFT and RFT datasets, there was an increase in the signal (i.e. a strong excess of mutated reads in NRFT). This can be seen in the bottom plot on the following page, where both curves (for the AB-PGT and VeriSeq PGS platforms) increase on the right side. This indicates that in families with a significant difference between NRFT and RFT embryos, this difference is reflected in the excess of mutated reads in NRFT compared to RFT.

### Supplementary Data 13 (additional 1)

VeriSeq PGS

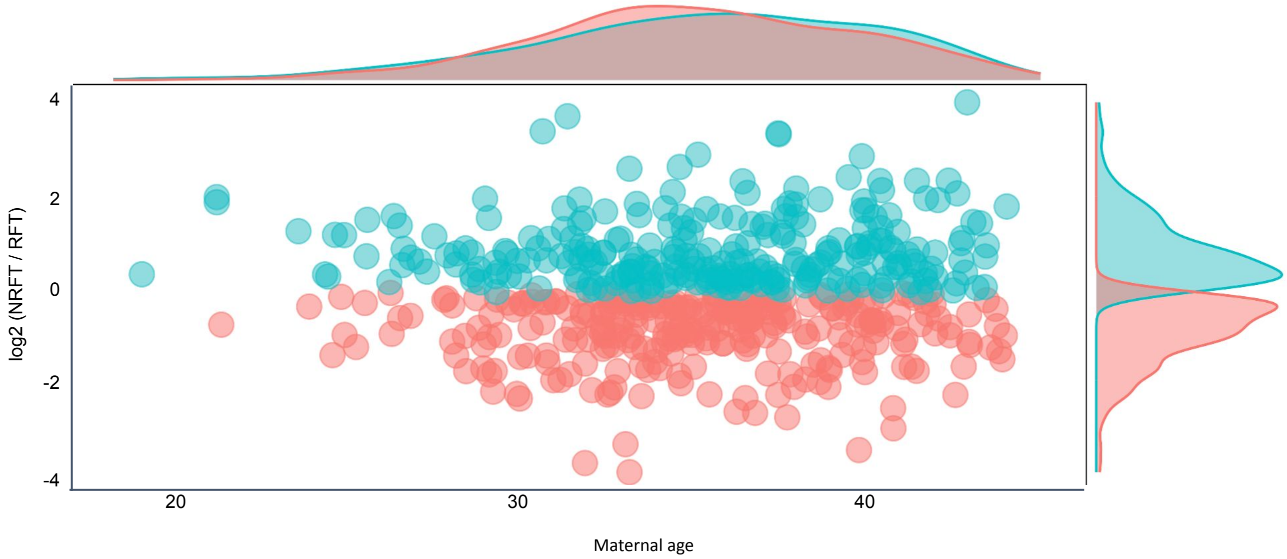


AB-PGT

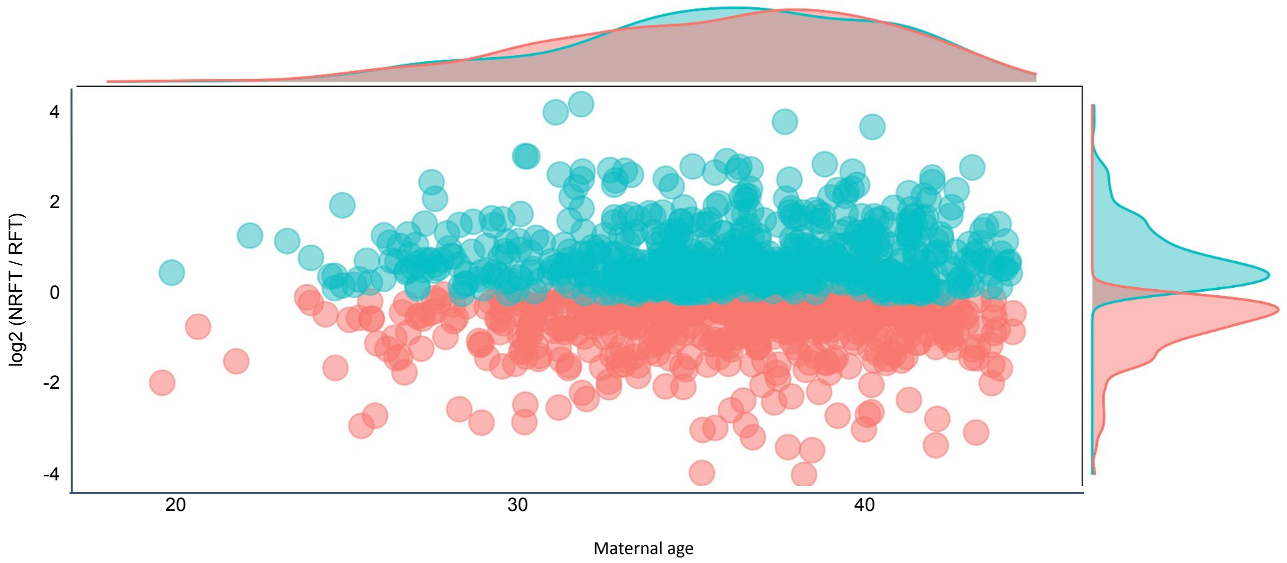


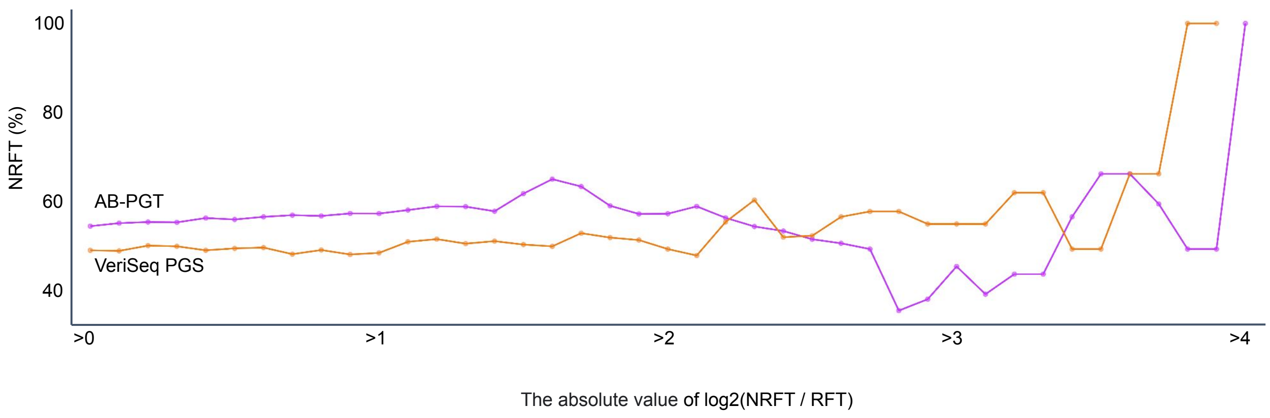

#### Supplementary Data 14

**Decision tree classifier confirms an especial importance of mtDNA content among young mothers.**

In order to reveal combined effects of age and mtDNA content on clinical outcome, we utilized Decision Tree Classifier, using it as sorting algorithm. This model, decision tree classifier, is based on conditional control statements, which binary and recursively split the data-set into subsets to maximize the information gain. In our case the classification rules should split embryos into NRFT (red crosses) and RFT (green circles) embryos based on two variables: mother's age and mtDNA content. An optimal decision tree is defined as a tree that accounts for most of the data, while minimizing the number of levels (questions). We used implementation from widespread Scikit-Learn python library in order to produce sorting rules, which distinguish RFT embryos from NRFT in a most efficient manner. For training of classifier we split dataset into train and test subsets, trained decision tree of maximum depth 2 and Shannon information gain as optimization criterion. While decision tree itself does not show quality often required by healthcare professionals in test dataset, decision boundaries we observe could serve as additional evidence for negative effect of excessive mtDNA content. Development of decision support algorithms with quality appropriate for clinical use most probably will require additional sources of information. AB-PGT data were used for the decision tree classifier as the most abundant dataset.

| | Precision | Recall | F1-score | Support |
|---|---|---|---|---|
| NRFT | 0.65 | 0.58 | 0.61 | 2795 |
| RFT | 0.57 | 0.63 | 0.6 | 2428 |
| Accuracy | - | - | 0.61 | 5223 |

Optimal boundaries acquired via decision tree

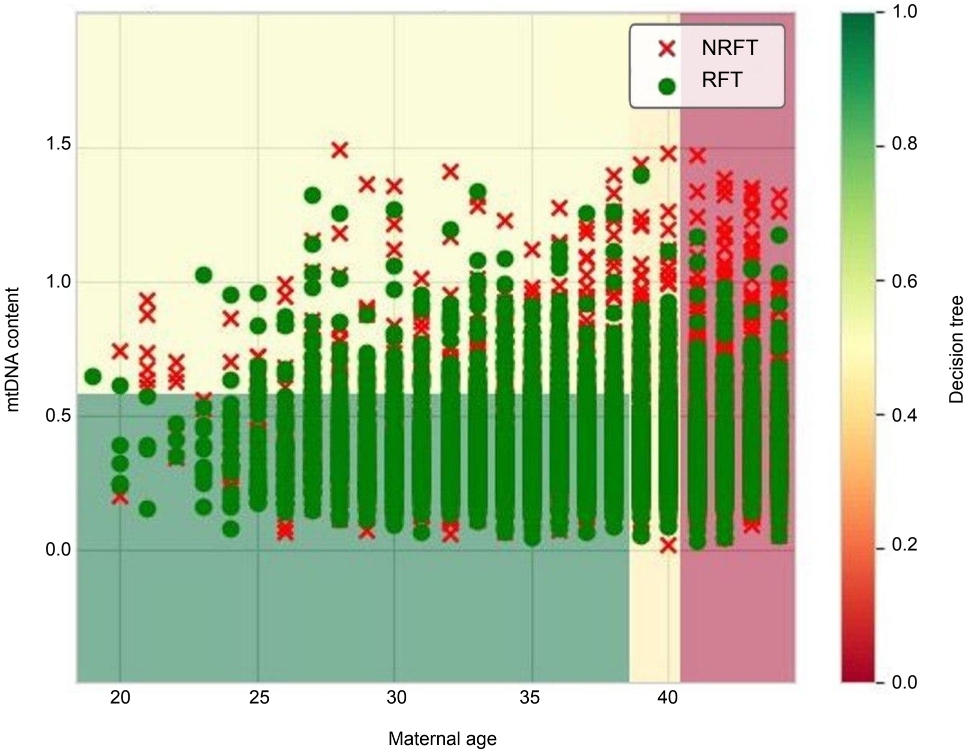

#### Supplementary Data 14 (additional 1)

**Decision tree classifier confirms an especial importance of mtDNA content among young mothers.**

First decision in the decision tree splits all samples according to the age: below or equal 38.5 or higher.

Among young mothers (<= 38.5) mtDNA content become the most important to split RFT against NRFT embryos: decreased mtDNA content (<= 0.595) is associated with maximal fraction of RFT (57.5%).

56.2% incidence of NRFT when the mtdna content is above 0.595 and the age is below 38.5.

The incidence of NRFT slightly increases to 58.3% for individuals with an age between 39 and 40 years, and further increases to 70.9% for individuals above 41 years of age.

AB-PGT data were used for the decision tree classifier as the most abundant dataset.

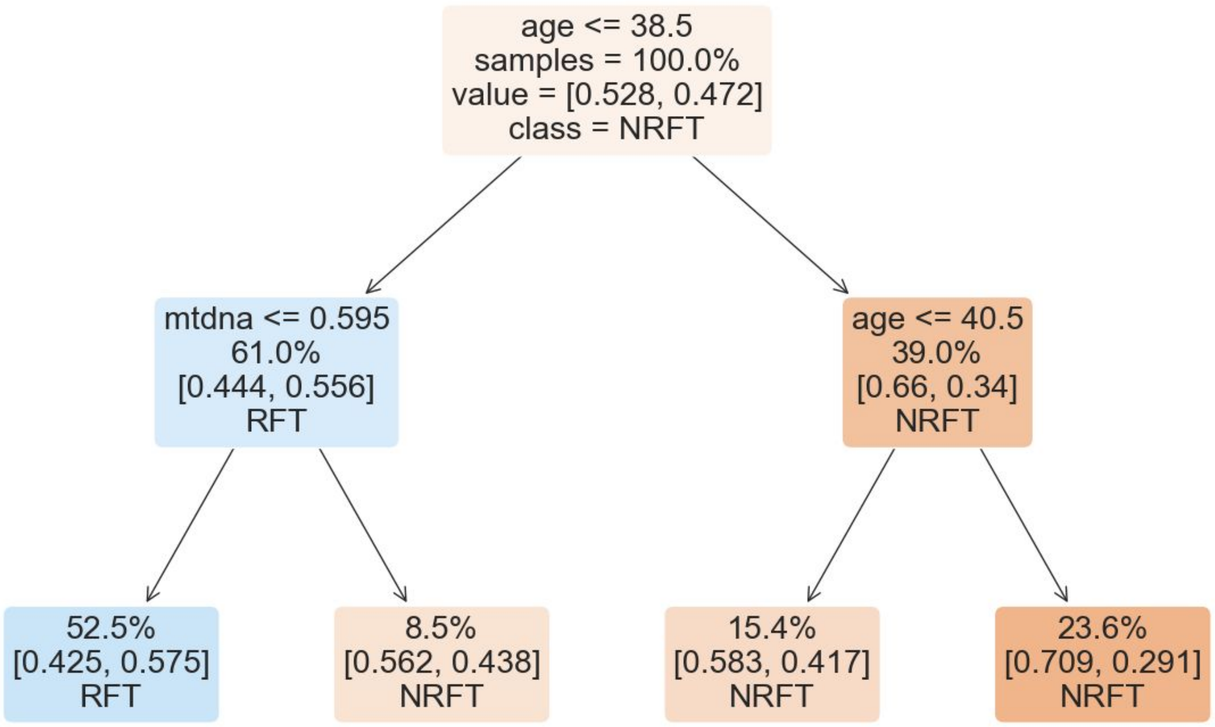
